# Domain-specific and domain-general signatures of dyscalculia in computational models of behavior and parietal cortex

**DOI:** 10.64898/2026.10.08.757623

**Authors:** Maike Renkert, Gilles de Hollander, Caroline Biegel, Sofie L. Valk, Karin Kucian, Christian C. Ruff

## Abstract

Developmental dyscalculia is a learning disorder characterized by marked difficulties in numerical cognition, yet its cognitive and neural underpinnings remain debated. Competing accounts attribute dyscalculia either to domain-specific imprecise representations of number or to domain-general deficits in the maintenance and integration of numerical information. Here, we test both accounts in 33 adolescents and young adults with dyscalculia and 33 age- and gender-matched controls (15-23 years) who performed a sequential numerosity comparison task during fMRI. Joint computational modeling of choices and response times allowed us to dissociate perceptual and memory noise from decision variables governing the speed-accuracy tradeoff, while complementary neuroimaging analyses characterized neural number representations and their network embedding. Specifically, we quantified the precision of parietal numerosity tuning with numerical population receptive field (nPRF) modeling and derived individualized functional network maps of the same region using community detection algorithms applied to connectivity patterns. Dyscalculic participants showed greater perceptual and memory noise but also a more cautious response criterion, trading speed for accuracy to compensate for representational imprecision. Neurally, the numerosity model explained less variance in parietal cortex, suggesting less precise numerical representations, and individualized network maps assigned less of this cortex to the dorsal attention network and more to a visual network. Rather than supporting a dichotomy between the two accounts, our findings suggest that dyscalculia reflects both less precise numerical representations in parietal cortex and an altered allocation of this cortex to distributed functional systems.

## Introduction

Developmental dyscalculia (DD) is a specific learning disorder affecting number and math skills that cannot be explained by low general intelligence or inadequate schooling, with an estimated prevalence of 3–6%^1,2^. DD impairs mental health^1,2^ and employment opportunities^3^, with broader societal consequences including costs to the national economy^4^. Despite this prevalence and impact, little is known about the etiology and precise symptomatology of DD, which is also reflected in the limited treatment options currently available. One factor complicating the development of homogeneous diagnostic criteria and treatments is the marked heterogeneity of the disorder^5,6^. Behaviorally, affected children can struggle with a wide range of numerical skills, such as counting, magnitude processing, transcoding between number words, digits and quantities, or spatial number representation^5,7^. But impairments are not confined to number skills: deficits in domain-general abilities, in particular working memory (WM) and attention, are frequently observed as well ^8,9^. It therefore remains unclear whether DD mainly arises from a domain-specific deficit in numerical representation or from a domain-general impairment in cognitive processes that support numerical reasoning^5,7^.

An influential domain-specific account proposes that DD originates from an imprecise approximate number system (ANS)^10–14^. This system, also referred to as the number sense, is thought to hold an abstract mental representation of non-symbolic numerosities that is independent of sensory modality, enabling rapid and effortless estimation and comparison of quantities without counting^15,16^. It is already present in infancy and may lay the foundation for symbolic number competencies and, ultimately, any type of mathematical thinking^15–18^. According to this number sense deficit theory, dyscalculic individuals have noisier internal representations of numerosity ^10,12^, although it remains debated whether such representational imprecision constitutes a primary cause or one component of a broader cognitive profile.

Domain-general theories, in contrast, propose that dyscalculia arises from impairments in processes that maintain, integrate, and manipulate numerical information over time, such as working memory, attentional control, and executive coordination. Several studies have highlighted visuospatial working memory as one prominent instantiation of such constraints^9,19^. One study^9^, for example, argued that poor numerical performance in dyscalculia relates mainly to deficits in visuospatial working memory, short-term memory, and inhibitory control rather than an imprecise number sense per se. Consistent with this view, visuospatial working memory capacity repeatedly predicts arithmetic abilities in both typical and atypical development^19,20^, and disruptions of working memory networks have been proposed as a core vulnerability factor in dyscalculia^21^. From this perspective, problems with numerical cognition stem less from an altered number sense than from limits on domain-general working memory resources that are crucial for both mathematical and visuospatial tasks.

At the neural level, DD has consistently been associated with structural and functional abnormalities in the intraparietal sulcus (IPS)^22,23^. This region, together with the adjacent postcentral sulcus, is central to numerical and mathematical cognition^24,25^; we therefore refer to it here as the numerical parietal cortex (NPC)^26^. Importantly, dyscalculia-specific alterations of the NPC (e.g.^27–32)^ are compatible with both accounts. The NPC is implicated in core numerical magnitude processing, such as the representation and comparison of quantities^26,33,34^, but also in broader cognitive operations, including the maintenance and manipulation of numerical information in working memory, attentional control, and the integration of symbolic and non-symbolic representations^35,36^. Abnormalities in this region could therefore affect both basic numerical processing and domain-general mechanisms supporting mathematical performance, and the previous literature cannot arbitrate between these views.

Consistent with the domain-specific account, evidence from neuroimaging and electrophysiology highlights the crucial role of the NPC in representing abstract numerical quantities^26,37,38^. This representation is modality-independent^39,40^ and extends to other kinds of magnitude such as time, space, or mass^33,41^. Neurons in parietal magnitude areas are tuned to specific numerosities, with peak responses at a preferred quantity that decrease monotonically with logarithmic distance from it^37,42^. These numerosity-tuned populations are topographically organized and can be visualized with numerical population receptive field (nPRF) modeling of fMRI data^26^. How well a presented number is encoded in, and can be decoded from, these nPRF maps relates to individual differences in behavioral numerical acuity^43–45^. nPRF modeling thus provides a handle on the fidelity of neural number representations, and we employ it here to test the domain-specific account.

With respect to more domain general perspectives, the IPS is also critically involved in visuospatial working memory and attentional control beyond its role in magnitude representation^35,36^. Working memory, however, is not implemented by a single cortical region but is widely considered an emergent function of dynamic interactions among distributed brain networks^35,46,47^. The influential sensory recruitment theory of working memory^48^ proposes that the sensory areas that process incoming information in a perceptual domain are also recruited to keep similar information “online”. Strikingly, the more abstract feature of numerosity appears to be stored in transmodal parietal cortex^49^, supporting the idea that this is where information is abstracted away from unisensory features, representing primarily the relational structure between numerical quantities^39,50^. Frontoparietal networks, in turn, are thought to control and gate access to this maintained activity through tight coupling and their own sustained activity^46,51^. Understanding potential working memory deficits for magnitude information in dyscalculia therefore requires investigating brain connectivity, in particular of the IPS and frontoparietal networks.

Indeed, previous studies have reported dyscalculia-related alterations in functional^52–54^, effective^55–57^, and structural^56,58,59^ connectivity. However, these findings stem from heterogeneous samples, age ranges, and methodological approaches, and often focus on isolated connections or predefined networks. It has therefore remained difficult to attribute connectivity differences to specific computational mechanisms, or to integrate them into a coherent account of how large-scale network organization constrains numerical processing in dyscalculia.

We therefore focus on one candidate neural substrate that maps conceptually well onto the domain-general account: the frontoparietal system in which the NPC is embedded. This network is crucial for solving arithmetic tasks^27–30^ but engages for a wide range of cognitive challenges, such as tests of fluid intelligence and non-verbal reasoning^60,61^. Given this breadth, it has been named the multiple-demand system, arguing that these regions support the assembly of structured mental programs, defining and controlling series of sub-operations, and are recruited more strongly as task complexity and working memory load increase^60,61^. Framing the same allocation of resources as a question of what is worth attending to at a given moment, others^62,63^ refer to a very similar set of regions as the dorsal attention network (DAN), the label we adopt here. Whatever the label, this network’s hypothesized contributions are precisely the cognitive operations that domain-general accounts of dyscalculia implicate.

Studying such a network in a group comparison is complicated by the fact that association cortex, to which frontoparietal DAN regions belong, aligns poorly across individuals in standard anatomical spaces^64,65^, because borders defined by macroanatomy, microstructure, and function diverge increasingly across the cortical hierarchy^66,67^. We therefore employed precision functional mapping to derive individualized network maps from functional connectivity data^68,69^, as such intrinsic patterns appear more informative about individual cortical organization than macroanatomy^70^. This allowed us to quantify individual and group differences in how much cortical area is devoted to the DAN, and to relate this measure to behavior. Cortical territory has been linked to behavior before: the extent and topology of frontoparietal networks predict executive function in children^64,65^ and cortical magnification in primary visual cortex predicts visual acuity^71^. We use this approach to investigate how DAN territory and topology, specifically in the parietal lobe, differ in dyscalculics, as a neural proxy for potential domain-general impairments that complements the nPRF measures targeting the domain-specific side.

Taken together, we here combine these neural analyses with computational modeling of behavior to characterize, in one sample and at both the behavioral and neural level, how basic numerical cognition, domain-general cognitive functions, and the underlying neural processes are altered in dyscalculia. Participants performed a sequential numerosity comparison task that places demands both on the perceptual approximation of non-symbolic numerosities and on the maintenance and comparison of this information across time. At the behavioral level, a computational cognitive model dissociates distinct sources of cognitive noise as well as evidence accumulation parameters, allowing us to separate perceptual precision from noise arising during memory and information integration, and from compensatory response selection strategies underlying speed-accuracy tradeoffs. At the neural level, we combine nPRF modeling as a direct, theory-driven measure of neural numerosity tuning^43–45^, with precision functional mapping, which quantifies how much of parietal cortex is devoted to the DAN, and thus relates to the maintenance and integration processes central to domain-general accounts. Beyond testing each measure for group differences, we finally ask whether these behavioral and neural measures placed on the same conceptual axis are quantitatively related to one another, and how separable the domain-specific and domain-general components actually are.

## Results

### Study population and experiment

We recruited 33 adolescents with dyscalculia and 33 age- and gender-matched controls (ages 15–23 years, 28 females in each group - see table 1). Dyscalculia diagnoses were made by a pediatric neuropsychologist at the children’s hospital. All participants completed a comprehensive assessment battery, including anatomical MRI, fMRI during cognitive tasks, standardized cognitive tests, and self-report questionnaires.

**Table 1.** Participant demographics and standardized test performance by group. IQ screener: ^72^; Corsi span: Corsi block-tapping task^73^, measure of visuospatial short-term memory; DEMAT: ^74^, standardized mathematical achievement test; Math anxiety & confidence: self composed questionnaires - inspired by MAI^75^; Panamath Weber fraction: ^76^, measure of approximate number system acuity. Group differences were assessed using 2-sample TTest or Mann–Whitney U tests depending on normality

| Measure | Control (mean ± SD) | Dyscalculia (mean ± SD) | statistics | p |
| --- | --- | --- | --- | --- |
| Gender (f / m) | 28 / 5 | 28 / 5 | - | - |
| Age (years) | 19.18 ± 2.86 | 19.09 ± 2.71 | U(33, 33)=553.50 | 0.912 |
| IQ screener (visuospatial reasoning) | 102.58 ± 15.87 | 93.79 ± 12.93 | t(64)=2.47 | 0.016 |
| IQ screener (verbal) | 107.27 ± 12.13 | 103.03 ± 12.05 | t(64)=1.43 | 0.159 |
| visuospatial short-term memory (Corsi span) | 7.48 ± 1.20 | 6.36 ± 0.86 | t(64)=4.36 | < 0.001 |
| Mathematical achievement (DEMAT test) | 33.30 ± 9.50 | 17.88 ± 8.93 | t(64)=6.80 | < 0.001 |
| Math anxiety | 2.34 ± 0.80 | 3.73 ± 0.89 | U(33, 33)=140.00 | < 0.001 |
| Math confidence | 3.15 ± 0.69 | 1.65 ± 0.81 | U(33, 33)=990.50 | < 0.001 |
| Panamath (Weber fraction) | 0.18 ± 0.06 | 0.21 ± 0.11 | U(33, 33)=429.00 | 0.140 |

Participant characteristics and standardized test performance are summarized in table 1. All participants had IQ scores within the normal range. The groups were broadly comparable (all statistics in table 1) in general intelligence as measured by verbal IQ, although the dyscalculia group showed slightly lower visuospatial reasoning (pattern recognition) IQ scores. As expected, participants with dyscalculia performed substantially worse on standardized mathematical achievement tests and reported higher levels of math anxiety and lower math confidence. In addition, dyscalculic participants showed reduced visuospatial short-term memory capacity, as measured by the Corsi block-tapping task. No group differences were observed in Panamath Weber fractions, indicating comparable basic numerosity acuity across groups.

### Behavioral task and cognitive modelling

To characterize the changes in neurocognitive magnitude processing in dyscalculia, we acquired behavioral and neural data for a perceptual magnitude comparison task, performed in the scanner (Fig.1A). On each trial, participants viewed two sequentially presented “coin clouds” and indicated which set contained more coins. The two displays were separated by a relatively extended, jittered interval of 6–9 seconds, which aided in the independent estimation of the haemodynamic responses evoked by each of the two stimulus presentations. Participants were allowed to respond as soon as the second display was presented.

**Figure 1.**
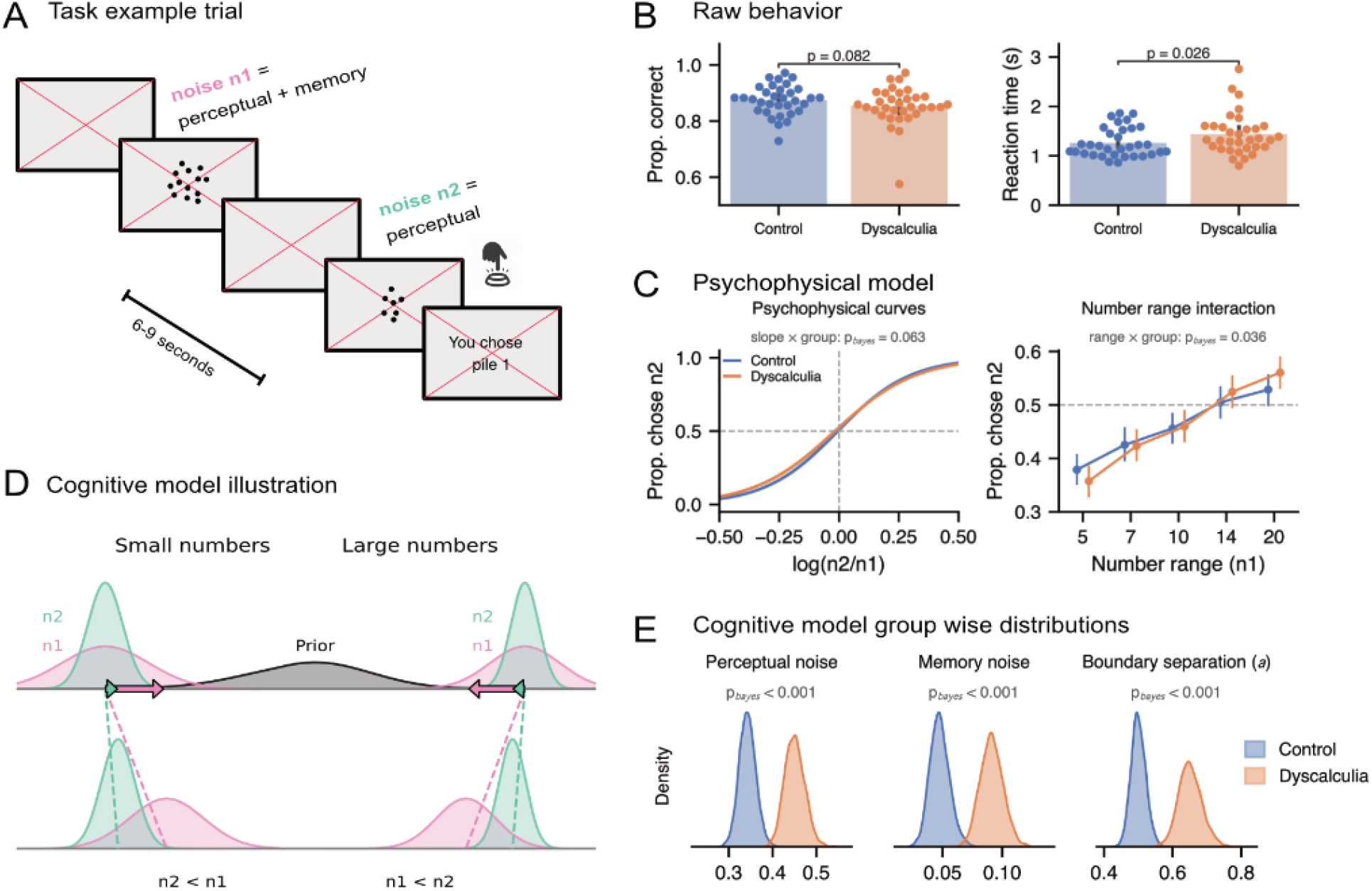
Non-symbolic magnitude comparison: task, behaviour, and psychophysical modelling. N = 66 (33 per group); 180 trials per participant. (**A**) Example trial illustrating the sequential presentation of the two non-symbolic numerosities as coin clouds, separated by a jittered interval (6–9 s). Participants indicated which array appeared more numerous to them upon presentation of the second set. (**B**) Raw behaviour. Proportion of correct choices and response times, one dot per participant; bars show the group mean and error bars the 95% bootstrap confidence interval of the mean. Groups were compared with a one-sided Mann–Whitney U test (because not normally distributed), testing the a-priori directional hypothesis that participants with dyscalculia are less accurate and slower; the p-value is given above each bracket. (**C**) Psychophysical modeling results. Aggregate behavior with proportion of choosing n2 to be larger as a function of log-ratio (n2/n1) and number range (if sets were rather small or large), with the posterior estimates of group effect sizes on these regressors as bayesian p-values indicated on top. (**D**) Illustration of the Bayesian perceptual memory choice model. Noisy likelihoods are formed for each numerosity display and combined with a prior over plausible magnitudes (grey) to yield posterior estimates that guide choice. Because the first-presented option (n1) must be retained across a delay, its likelihood is broader (pink) than that of the second-presented option (n2; green), reflecting additional memory-related noise. For small numerosities, likelihoods fall to the left of the prior and are therefore overestimated (central tendency), whereas the opposite occurs for large numerosities. These biases are amplified for noisier representations (e.g., n1 which also comprises memory noise), leading to a reverse choice bias: for small numbers, subjects are more likely to report n1 as larger, but for large numbers, they are more likely to report n2 as larger. (**E**) Group-level parameter estimates of the PMC-RDM model. Each density is the posterior distribution over the group-level mean of a parameter, perceptual noise, memory noise, and boundary separation (a), shown separately for each group on the softplus-transformed (positive) scale on which it enters the model. Above each panel, the bayesian p-value is the posterior tail probability of the group difference (dyscalculia − control).

Initial analyses of overall task performance confirmed that, as expected, adolescents with dyscalculia were slower (response time: Control = 1.26 ± 0.30 s, Dyscalculia = 1.44 ± 0.42 s, Mann–Whitney U (33, 33) = 392, one-sided p = 0.026) and accuracy was lower, in the predicted direction, though with weaker support (Control = 0.87 ± 0.05, Dyscalculia = 0.85 ± 0.07, U (33, 33) = 653.5, one-sided p = 0.082; Fig.1B). To capture more subtle differences in numerical decision-making in a principled manner, we fitted an integrated psychophysical (probit) model estimating the probability of reporting the second set (n2) as more numerous as a function of the numerical ratio between the two sets. The model included group (control vs. dyscalculia), number range (continuous predictor reflecting whether comparisons involved smaller or larger numerosities ^77^), and their interaction terms. Model fits revealed the expected shallower slope for DD but without reaching significance (slope x group interaction: mean = −0.58, 95% CI = [−1.31, 0.16], *p_bayesian_* = 0.063), consistent with the qualitatively similar psychometric functions at the aggregate level (Fig.1C, left). However, within both groups there was a clear main effect of the number range (mean = 0.034, 95% CI = [0.024, 0.045]), indicating an increased tendency to judge the second display (n2) as more numerous when comparisons involved larger numerosities in general (Fig.1C, right; note that our experimental design made sure that for each *n1*, we used the same set of *fractions*, making the range of ratios identical). This bias was more pronounced in dyscalculic participants (number range × group interaction: mean = 0.013, 95% CI = [−0.001, 0.028], *p_bayesian_* = 0.036; Fig.1C, right). Such a range-dependent bias likely arises because the first stimulus, unlike the second, must be retained in memory across the delay and is therefore represented more noisily; when this degraded estimate is combined with prior expectations about the stimulus distribution, it is drawn toward the prior’s mean ^78^. Since the weight given to the prior scales with this mnemonic uncertainty, the stronger effect in dyscalculic participants points to a noisier working memory system, greater reliance on the prior, or both ^79^.

To understand the specific cognitive alterations underlying the observed differences, we implemented an extended, dynamic Bayesian perceptual choice model (PMC; illustrated in Fig.1D), originally developed in earlier work by our group ^45,80^. This model enables the dissociation of perceptual noise, reflecting the precision with which numerosity is extracted from an acute visual display, from memory-related noise arising during the retention of the first display across the inter-stimulus interval, directly addressing our primary interest in distinguishing domain-specific from domain-general sources of numerical imprecision. On each trial, the two numerical displays were treated as incoming evidence combined with a prior expectation of plausible magnitudes to form a posterior estimate—the percept that guided the final choice. Importantly, because the displays were presented sequentially, the precision of the first option (n1) reflected both perceptual and memory noise, whereas the second option (n2) was influenced only by perceptual noise (i.e., the precision of the number sense). Note that this asymmetry can naturally explain the number-range effect: the noisier n1 estimate is shrunk more strongly toward the prior, so n1 is relatively overestimated at small numerosities and underestimated at large ones, biasing choices toward n2 in the larger ranges.

When comparing populations with different performance, one critical aspect of perceptual decision-making that must be taken into account is the speed–accuracy tradeoff. How much evidence a participant requires before committing to a response can mimic, or mask, a difference in the precision of the underlying representations ^81–83^. Our DD participants did respond slower while being only somewhat less accurate, a combination that could reflect greater response caution rather than - or in addition to - noisier representations^81–83^. We therefore extended the PMC model with a racing diffusion likelihood^84^ under the advantage parameterisation of van Ravenzwaaij et al.^85^, so that it jointly predicts choice proportions and response times (see methods for further discussion of why we use a race model instead of standard diffusion model). This PMC-RDM model thus adds four free parameters: a baseline drift *w*_0_, which adds to both accumulators regardless of the stimuli and is interpretable as response urgency; a weight *w_sum_* on the summed magnitude, capturing the magnitude effect on response times; a decision threshold *a*, the evidence each accumulator must reach, indexing response caution; and a non-decision time *t*_0_, absorbing sensory and motor delays. Critically, with the PMC-RDM, the precision of percept formation (sensory noise) can be separated from the decision-related parameters that would otherwise mask it, chief among them the response caution captured by the threshold *a*.

Model parameters were estimated within a hierarchical Bayesian framework, allowing group membership to modulate all parameters. Parameter estimates of the PMC-RDM showed that both perceptual and memory processes are impaired in DD (Fig.1E). Memory-related noise was estimated to be higher in dyscalculic participants (mean difference = 0.044, 95% CI [0.019, 0.069], *p _bayesian_* = 0.001; dyscalculia: mean = 0.091, 95% CI [0.073, 0.110]; control: mean = 0.047, 95% CI [0.033, 0.065]). Likewise, perceptual noise was also higher in the dyscalculia group (mean difference = 0.108, 95% CI [0.051, 0.167], *p_bayesian_* < 0.001; dyscalculia: mean = 0.451, 95% CI [0.409, 0.497]; control: mean = 0.343, 95% CI [0.307, 0.381]). Of the additional parameters introduced by the race diffusion choice function, boundary separation *a* was also estimated to be higher in dyscalculic participants (mean difference = 0.151, 95% CI [0.077, 0.232], *p_bayesian_* < 0.001; dyscalculia: mean = 0.651, 95% CI [0.588, 0.722]; control: mean = 0.500, 95% CI [0.460, 0.544]). This is consistent with a more cautious response criterion in dyscalculia, and fits with the somewhat slower response times observed at the raw performance level. Non-decision time t₀, by contrast, was not different between the groups (mean difference = −0.019, 95% CI [−0.082, 0.042], *p_bayesian_* = 0.274; dyscalculia: mean = 0.363, 95% CI [0.317, 0.410]; control: mean = 0.382, 95% CI [0.339, 0.425]), as would be expected if the groups do not differ in basic sensory and motor delays. Baseline drift *w*_0_ was likewise somewhat elevated in dyscalculic participants (mean difference = 0.086, 95% CI [0.016, 0.159], *p_bayesian_* = 0.009; dyscalculia: mean = 0.398, 95% CI [0.345, 0.458]; control: mean = 0.312, 95% CI [0.267, 0.361]). We interpret this elevated urgency as a downstream consequence of the greater response caution in dyscalculic participants: because their slower responses more frequently approached the response deadline, they were under greater effective time pressure. Such deadline-induced time pressure has been shown to increase urgency rather than lower the decision threshold ^86^. Consistent with this account, dyscalculic participants missed the response deadline more often than controls (2.2% vs. 1.3% of trials; U = 679, p = 0.04).

To understand how much of these effects are depending on also taking response times and hence the decision process into account, we also estimated the model in its choice-only informed version. The memory-noise difference proved resilient to dropping the RDM back-end, and was estimated in the same direction and at a comparable magnitude (mean difference = 0.042, 95% CI [−0.003, 0.087], *p_bayesian_* = 0.032; control: mean = 0.122, 95% CI [0.09, 0.155]; dyscalculia: mean = 0.164, 95% CI [0.134, 0.193]). Conversely, the perceptual-noise difference did not persist: there was little indication of a group difference here, with the point estimate if anything running in the opposite direction (mean difference = −0.016, 95% CI [−0.051, 0.019], *p_bayesian_* = 0.188; control: mean = 0.103, 95% CI [0.078, 0.129]; dyscalculia: mean = 0.087, 95% CI [0.063, 0.112]). Taken on its own, this model variant would thus locate the group difference only in the maintenance of numerical information across the inter-stimulus interval, which one would expect if elevated response caution goes unmodelled: a choice-only model can attribute accuracy only to representational precision, so the accuracy bought by a wider threshold is absorbed into the perceptual noise estimate. To ensure that this interpretation is accurate, we simulated 1,000 datasets from the fitted PMC-RDM posterior, where the DD group had by construction both higher perceptual noise and a wider threshold, and refitted the choice-only PMC. Also from these simulated data, it recovered *lower* perceptual noise in the simulated DD group in 82% of fits (mean difference −0.15, 95% CI [−0.16, −0.14]), closely reproducing the reversal seen in the real data (−0.19). This also showed that memory noise escapes misidentification for response caution because it is identified by the contraction bias, a pattern in how choices depend on number range rather than overall accuracy which caution does not mimic. Thus, our analyses and simulations confirmed that our dyscalculic participants do have elevated perceptual and memory noise, but that the former is masked by the greater response caution they apply potentially as a compensatory strategy. Correlations across participants between the three levels of description of the task (raw performance, the probit model, and the PMC-RDM) confirmed that the model parameters captured what they were meant to capture: for example, perceptual noise was closely related to the probit slope, memory noise to the number-range effect, and boundary separation to response time (Supplementary Figure 1 and Supplementary Note 1).

Taken together, these analyses suggest that the two (domain-specific vs domain-general) putative components of cognitive differences of dyscalculia are both expressed but not equally easy to detect. More domain-general differences appear across several independent descriptions: in the standardized tests of visuospatial reasoning and short-term memory, in the number-range effect of the probit model, and in the memory noise parameter (both with and without modeling the decision process with the RDM). The domain-specific difference does not directly show up in coarser indices (overall accuracy, psychometric slope, and Panamath Weber fractions), but becomes apparent once response times are modelled alongside choices, a description that separates representational noise from the elevated response caution that otherwise conceals it.

### Neural modelling

To examine the brain basis of numerical processing deficits and general cognitive deficits in developmental dyscalculia, we focused on the numerical parietal cortex (NPC), a region repeatedly implicated in both numerical cognition and dyscalculia (e.g ^27–32^) and applied two complementary neural modelling approaches that target distinct properties of NPC function. First, we assessed the *precision of numerical representations* using numerical population receptive field (nPRF) modelling, which provides a neural proxy for number sense acuity. Second, we characterized *large-scale functional integration* of the NPC using precision functional mapping-based connectivity analyses, capturing how this region is embedded within broader cortical networks. Lastly, we also investigated the spatial topology of *individual maps of the dorsal attention network (DAN)*, also referred to as the multiple-demand system ^60^, due to its domain general role in maintaining and manipulating task-relevant information.

### Numerical representations in the NPC

As a first step, we examined whether dyscalculia is associated with altered neural precision of numerical representations in the NPC. To this end, we employed numerical population receptive field (nPRF) modelling, an encoding–decoding framework that quantifies population-level tuning to numerosity and has been linked to behavioral measures of numerical acuity ^43–45^. We fitted numerical receptive field models to each voxel’s single-trial activity at the time point of presentation of the first numerosity stimulus (n1). The model assumes that each voxel pools over a population of numerosity-tuned neurons and therefore responds most strongly to one preferred numerosity, with responses falling off for numerosities increasingly distant from it on a logarithmic scale. Each voxel’s tuning is described by four parameters: its preferred numerosity, the width of its tuning (dispersion), and the amplitude and baseline of its response (see Methods for the formal specification). Following previous work ^43–45^, the spatial extent of the group-average variance-explained map was used to derive a functionally defined NPC mask, which serves all analyses reported in this section.

First, we compared across groups how well neural responses in this region were explained by a model that characterises numerosity tuning with a log-Gaussian function. Quantified as the mean variance explained (R2) across NPC vertices from the encoding model, this measure was lower in dyscalculic participants than in controls (control: 0.0233 ± 0.0069, 95% CI [0.0208, 0.0257]; dyscalculia: 0.0198 ± 0.0064, 95% CI [0.0175, 0.0220]; U = 705.0, p = 0.021; Fig.2B). Given the modest sample size (see Methods for a sensitivity analysis), we also tested whether any single participant drove this and the following neural group differences by repeating each test with one participant left out at a time. The difference in mean R² corresponded to a medium effect (d = 0.53, 95% CI [0.02, 1.07]), smaller than the effect size the design could reliably detect (see Methods), but remained below p = 0.05 in all leave-one-out samples (maximum p = 0.031). Two related measures showed the same difference, median R2 (control: 0.0195 ± 0.0052, 95% CI [0.0177, 0.0214]; dyscalculia: 0.0168 ± 0.0044, 95% CI [0.0153, 0.0184]; p= 0.013) and amount of vertices with R2 > 0.02 (control: 665 ± 245, 95% CI [577, 753]; dyscalculia: 514 ± 249, 95% CI [425, 602]; p = 0.016; Fig.2B). These results were robust across a range of different spatial thresholds (number of vertices) and amount of area included (mean R2), and held irrespective of whether the measures were derived in surface or individual volumetric (native T1) space (all stats for these control analyses are given in Supplementary Table 1).

**Figure 2.**
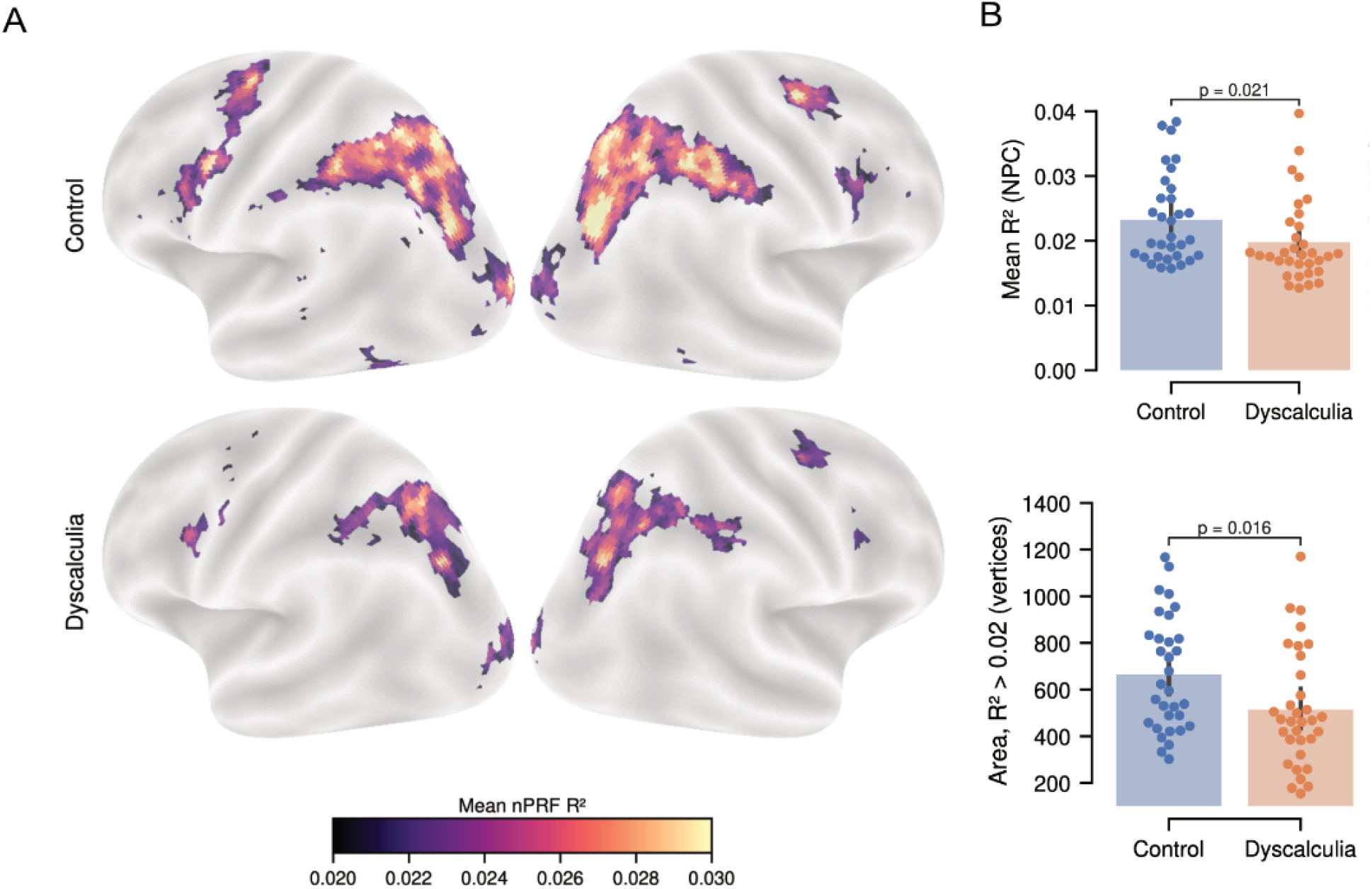
Numerical coding in the parietal cortex. N = 65 (32 controls, 33 participants with dyscalculia). (**A**) Group-average nPRF variance explained R2 maps, thresholded at 0.02 (beginning of the scale) shown separately for controls (top) and dyscalculic participants (bottom) on the inflated *fsaverage5* surface. Both groups show the expected parietal focus, with lower and less spatially extended values in dyscalculia (the NPC mask used for all analyses reported here was derived from the corresponding average across both groups combined). Examples of individual (thresholded) nPRF maps are shown in supplementary figure 3. (**B**) Group comparisons of two summaries of the within-mask R² distribution: mean R² and the number of vertices exceeding R² = 0.02. Each dot is one participant; bars show the group mean and error bars the 95% bootstrap confidence interval of the mean. Groups were compared with a Mann–Whitney U test for mean R² and an independent-samples t-test for the vertex count (non-parametric tests were used where the measure was not normally distributed; see Methods), two-sided; the p-value is given above each bracket.

Secondly, as done in previous studies with a classical encoding-decoding framework ^43–45^, we investigated how well the presented number could be decoded from the neural data given the model fit: Inverting the encoding model together with an additional noise model yields a decoded numerosity for each held-out trial, and its correlation with the presented numerosity indexes the individual precision measure (see Methods). In contrast to R2 measures that only relied on the encoding model, this measure was similarly distributed in both groups (control: 0.163 ± 0.142; dyscalculia: 0.168 ± 0.117; mean difference = −0.005, 95% CI [−0.068, 0.058], t = −0.15,p = 0.88), even though decoding accuracy was correlated with mean R2 measures (Spearman ρ = 0.65, p < 0.0001). Taken together, the nPRF analysis suggested that for dyscalculics, neural number representations in the NPC are less precise, but encoding model-based measures seem to be more sensitive to detect this difference than a decoding approach.

### Functional network composition of the NPC

To test the alternative perspective that the NPC may be differently embedded in large-scale cortical networks, we next quantified how much of its territory is assigned to different functional networks in each participant’s individual partition ^68,69^. Connectivity-based parcellation approaches derive such network allocations by grouping cortical nodes into communities (modules) with similar connectivity patterns, given that they represent a primary organizing feature of cortical function beyond anatomy ^70^. Precision functional mapping (PFM) applies the procedure to the connectivity matrices of individual subjects to derive individualized network maps, using the Infomap algorithm ^87^ for community detection. This is particularly relevant for association regions such as the NPC, whose topography varies substantially across individuals ^64,65,68^. With these individualized network solutions, we investigated (1) how cortical territory within the NPC region is functionally distributed and (2) how dorsal-attention network topology might differ between groups over the whole brain.

To label the individual maps consistently across participants, we first generated an all-subject average task network map (Fig.3A) from the average connectivity matrix, using the Cole-Anticevic network map ^62^ as a reference for label assignment, which in turn served as the reference for labelling the individual network maps. Of the canonical networks represented within the NPC, only three were present in every participant: dorsal attention (67.62 ± 13.49 cm²), somatomotor (13.17 ± 7.42 cm²), and visual-2 (10.33 ± 8.89 cm²). The remaining networks were only sparsely found in the NPC mask across subjects and were not considered further. Two of these three differed between groups: The NPC surface area assigned to the DAN was smaller in dyscalculic participants (control: 72.29 ± 12.96 cm²; dyscalculia: 62.94 ± 12.52 cm²; U = 761.00, p = 0.006), while that assigned to the visual-2 network was larger (control: 6.72 ± 6.33 cm²; dyscalculia: 13.94 ± 9.67 cm²; U = 295.00, p = 0.001). Both differences were sizeable (DAN: d = 0.73, 95% CI [0.27, 1.27]; visual-2: d = −0.88, 95% CI [−1.44, −0.42]), at or above the effect size the design could reliably detect (see Methods), and held when any single participant was left out (maximum p = 0.009 and 0.002, respectively). Somatomotor area, by contrast, was closely matched (control: 12.95 ± 7.81 cm²; dyscalculia: 13.39 ± 7.13 cm²; U = 517.00, p = 0.729). This pattern indicated a shift in the relative cortical territory devoted to functional systems with distinct connectivity profiles in DD, from a transmodal, attention-related system towards a unimodal, visual one (Fig.3B and E). These differences appeared to be regionally circumscribed: Whole-brain comparisons of network sizes revealed no group differences that survived correction for multiple comparisons (see Supplementary Table 2 for test statistics of all comparisons), indicating that the observed effects are unlikely to reflect global differences in network size.

**Figure 3.**
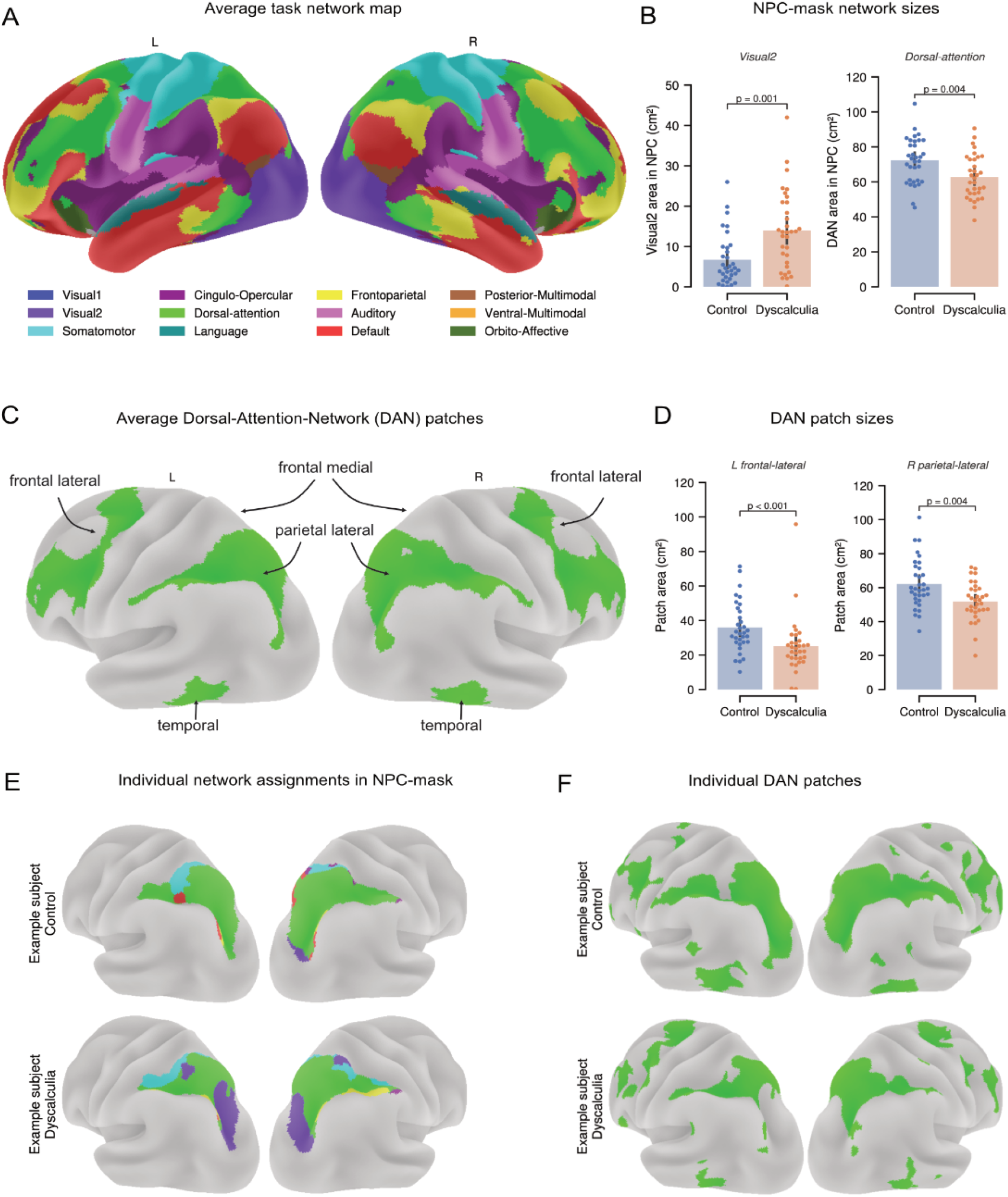
Connectivity analysis based on individual network maps. N = 66 (33 controls, 33 participants with dyscalculia). (**A**) Group-average task network map from precision functional mapping, shown on the partly-inflated *fsLR* surface (lateral views only). Network labels follow the Cole-Anticevic atlas. (**B**) Group comparison of visual-2 and dorsal attention surface area (cm²) within the NPC mask (control n = 33, dyscalculia n = 33). Each dot is one participant; bars show the group mean and error bars the 95% bootstrap confidence interval of the mean, p-value is uncorrected. See results methods for details on the statistical test. (**C**) Reference DAN patches derived from the group-average network map by connected-components analysis, with approximate anatomical labels (frontal-lateral, frontal-medial, parietal-lateral, temporal) assigned from patch centroid coordinates. Only lateral views are shown, so the frontal-medial patch is not itself visible - its position is indicated by the arrow. (**D**) Group comparison of individual DAN patch area (cm²) for the two patches that survived FDR correction across the eight bilateral patches. Conventions otherwise as in B, except that group sizes vary by patch (control 30–33, dyscalculia 28–33), since not every reference patch was recoverable in every participant’s individual network partition. All eight patches are reported in Supplementary Table 3. (**E**) Network assignments within the NPC mask for one example control (top) and one example dyscalculic participant (bottom). Only networks occurring within the mask are shown: dorsal attention (green), visual-2 (purple), somatomotor (cyan), cingulo-opercular, frontoparietal, and default. The two participants illustrate the range of NPC network composition across individuals. (**F**)Individual DAN partitions for the same two example participants, illustrating how patch topography varies across individuals. More example maps (with nPRF map overlays) are shown in supplementary figure 3.

Because the DAN extends well beyond the NPC, and is the network that supports the maintenance and manipulation of task-relevant information across a wide range of tasks^60^, we followed up on the NPC-focused analysis with an analysis at the level of the whole network. In each participant’s own whole-brain network parcellation, we decomposed the DAN territory into spatially contiguous patches, matched to reference patches from the group-average map (Fig.3C) by centroid position, and compared these between groups across the eight main bilateral patches. As expected, the two parietal-lateral patches were both smaller in dyscalculia, but only the right survived correction over the eight comparisons (U = 769.00, p = 0.004; p(FDR) = 0.016, Fig.3D; right p: U = 718.00, p = 0.027, p(FDR) = 0.071). Group means and all eight comparisons are given in Supplementary Table 3. Interestingly, we also found a group difference outside the parietal cortex, in the left frontal-lateral patch (U= 804.00, p < 0.001, p(FDR) = 0.002, Fig.3D left). Both patch differences that survived correction were stable when any single participant was left out (right parietal-lateral: maximum uncorrected p = 0.006; left frontal-lateral: maximum uncorrected p < 0.001). As expected, the right parietal-lateral effect largely recapitulates the DAN NPC-mask result, since the two measures are nearly redundant (r = 0.85, p < 0.001). Notably, DAN area within the NPC was also weakly associated with the mean R² nPRF measure (ρ = 0.30, p = 0.017; with group partialled out: ρ = 0.22, p = 0.083). While this may be expected, given that the two measures are derived from overlapping cortex (see Supplementary Figure 3 for example overlay maps from both neural models), it also means that they should not be treated as fully separable effects.

Taken together, our neural network modelling results suggest that the NPC is embedded differently in the brain’s functional networks in dyscalculia: Less cortical territory is devoted to dorsal attention network functions, and correspondingly more to visual network functions. Thus, in dyscalculia, the same stretch of parietal cortex bears more unimodal, sensory connectivity patterns at the cost of less transmodal, attention-related ones. This shift does not follow from globally different network sizes, but still is not totally confined to the NPC, since a left frontal-lateral DAN patch was also smaller in dyscalculic participants. The NPC therefore is both less responsive to numerosity (the domain-specific side) and differently functionally connected with the large-scale systems that support maintenance and integration of numerical information (the domain-general side); a finding that potentially mirrors the conclusions of the behavioral findings described before. The positive correlation between the model nPRF and PFM measures further suggests that these deficits are at least partly related.

### Continuous brain-behavior associations

On both the behavioral and neural level, we found multiple DD-specific alterations in measures that conceptually relate to cognitive components considered more domain-specific or domain-general, respectively. To investigate how closely these components correspond to each other across the behavioral and neural level, we computed partial Spearman correlations with group entered as a covariate, so that coefficients reflect covariation beyond the diagnostic split (see Methods). The full correlation matrix is shown in Fig. 4A. Two associations stand out, one on either side of the domain-specific/domain-general divide. On the domain-specific side, PMC-RDM perceptual noise was negatively related to mean nPRF R² (ρ = −0.42, p < 0.001; Fig. 4B), so that participants whose choices implied noisier perceptual encoding had less NPC territory where activity was well accounted for by the numerosity model. This was the only association that survived FDR correction across the full matrix (p(FDR) = 0.026). On the domain-general side, visuospatial IQ was positively related to DAN area within the NPC (ρ = 0.37, p = 0.003; Fig. 4C), such that participants with higher visuospatial IQ tended to have a larger NPC DAN territory. This association did not survive correction across the matrix (p(FDR) = 0.067), and we therefore consider it tentative. Relative to the sensitivity of the design (see Methods), the association with perceptual noise lay above the reliably detectable size and that with visuospatial IQ at its edge. When one participant was left out at a time, both associations remained of similar size (ρ ranges −0.48 to −0.40 and 0.32 to 0.43, respectively, all p < 0.05). After correction across the full matrix, however, the association with perceptual noise held in 65 of 66 leave-one-out samples, and that with visuospatial IQ in only 12 of 66, consistent with its tentative status. Because both neural measures (nPRF R² and NPC DAN area) are drawn from the same cortex (see Supplementary Figure 3 for examples of how nPRF and DAN maps overlap) and are weakly correlated (statistics noted above), we also asked whether the two links are separable or two views of one association (Williams tests for dependent correlations, one-tailed). Perceptual noise was more closely related to nPRF R² than to DAN area (difference in ρ = −0.45, 95% CI [−0.75, −0.15]; t(62) = −3.10, p = 0.001). For visuospatial IQ, the difference pointed in the expected direction, with a closer relation to DAN area than to nPRF R², but was less clear (difference = 0.25, 95% CI [−0.07, 0.54]; t(62) = 1.67, p = 0.051). The domain-specific link thus appears specific to its own neural counterpart, whereas the evidence for specificity on the domain-general side is weaker. Notably, the PMC-RDM memory noise parameter showed little association with DAN area (ρ = −0.05, p = 0.71), suggesting that the cognitive component related to the domain-general neural measure was mainly indexed by visuospatial IQ. Neither visual-2 area nor the left frontal-lateral patch, both of which showed group differences, was associated with any behavioral measure (all |ρ| ≤ 0.22, smallest uncorrected p = 0.079).

**Figure 4.**
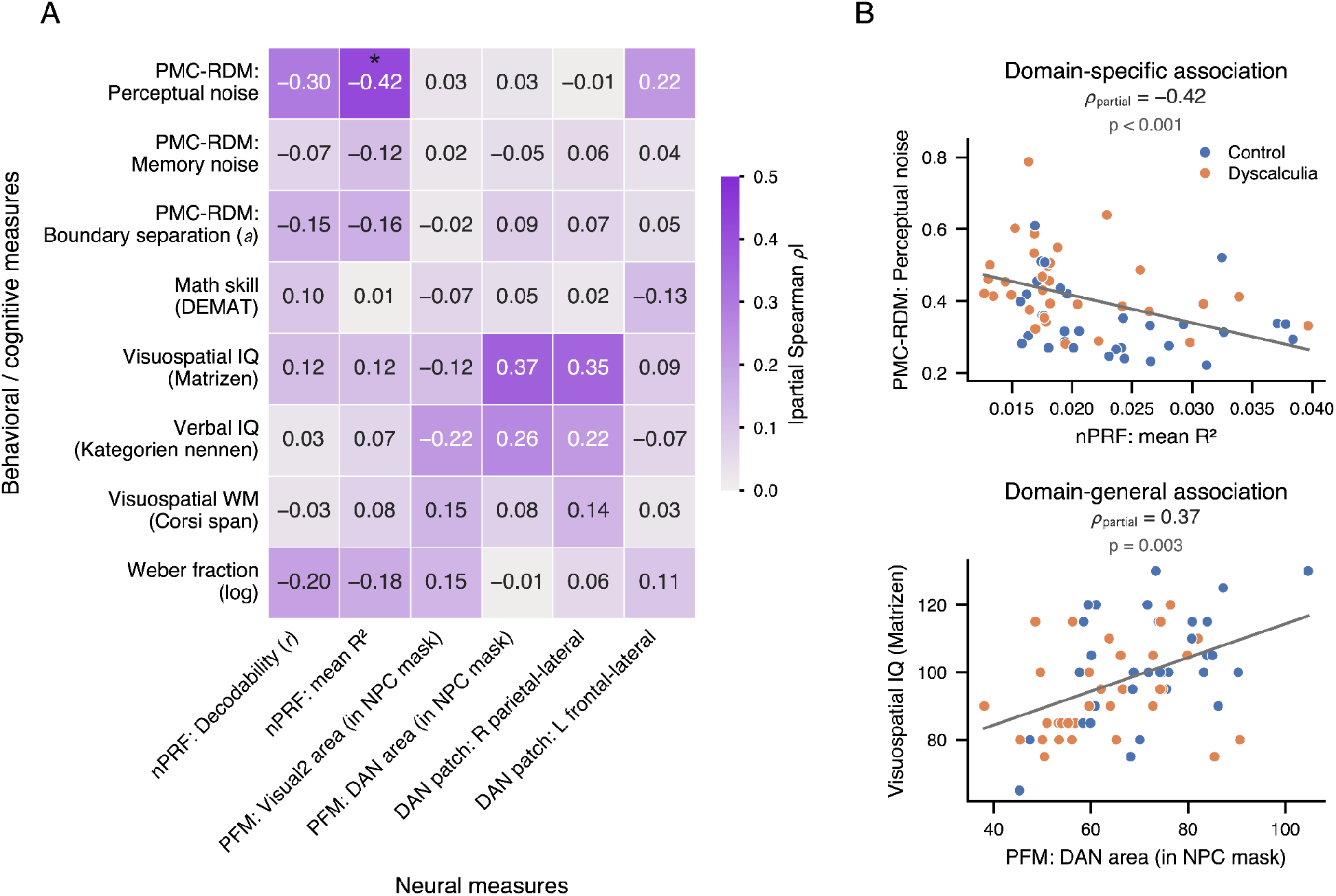
Brain-behavior correlations. N = 66 (33 controls, 33 participants with dyscalculia); n = 65–66 per cell with pairwise deletion, as one participant was excluded from the nPRF analyses (see Methods). (**A**) All associations between behavioral/cognitive and neural measures, with group partialled out. Each cell gives the partial Spearman correlation between one behavioral measure (rows) and one neural measure (columns), with group (dyscalculia vs. control) entered as a covariate, so that a coefficient reflects covariation beyond what both measures share with the diagnostic split. Printed values keep their sign; cell shading encodes |ρ| on a common 0–0.5 scale. Asterisks mark cells surviving correction for multiple comparisons across the full matrix (Benjamini–Hochberg FDR, * p < 0.05). Behavioral measures comprise the three PMC-RDM parameters (perceptual noise, memory noise, boundary separation a) and the standardized cognitive tests; neural measures comprise the two nPRF read-outs, the two networks most present within the NPC mask, and the two DAN patches that differed between groups. (**B, C**) The two associations framing the domain-specific/domain-general divide: PMC-RDM perceptual noise against mean nPRF R² (B; n = 65), which survived FDR correction across the matrix (p(FDR) = 0.026), and visuospatial IQ against dorsal attention area within the NPC (C; n = 66), which did not (p(FDR) = 0.067). Above each panel, ρ_partial is the group-partialled Spearman correlation from A and the p-value is uncorrected. Axes show raw measure values, not residuals; each dot is one participant, coloured by group; one participant with high perceptual noise lies outside the plotted range in B but is included in all statistics. The grey line is an ordinary least-squares fit through the pooled sample, shown as a visual aid only.

### Summary

Taken together, we found evidence for both accounts, at both the behavioral and the neural level, with different strength across measures. The clearest evidence came from the behavioral model, in which dyscalculic participants showed higher perceptual noise, the domain-specific component, and higher memory noise, the domain-general component, together with greater response caution that partly masked the perceptual difference. Neurally, the most robust differences concerned the network composition of the NPC, of which less was assigned to the dorsal attention network and correspondingly more to the visual-2 network. The numerosity model also explained less variance in the NPC of dyscalculic participants, a smaller difference that should be considered more tentative. Across participants, perceptual noise covaried robustly with nPRF R², and, more tentatively, visuospatial reasoning with NPC DAN area.

## Discussion

In this study, we tested, at both the behavioral and the neural level, two competing accounts of what differs between DD and neurotypical peers: a domain-specific account focusing on the precision of numerical representations, and a domain-general account focusing on the maintenance and integration of this information over time. We found evidence for both. Behaviorally, domain-general differences appeared across several independent measures: standardized tests of visuospatial reasoning and short-term memory, a number-range effect on choice that likely reflects increased working memory noise, and the memory noise parameter of the PMC-RDM. Domain-specific differences, in contrast, were not evident in coarser measures of number sense acuity (Panamath Weber fraction, choice accuracy, and psychometric slope), but became apparent in a cognitive model accounting for both choices and response times, as higher perceptual noise alongside a higher decision threshold that partly counteracted it. Neurally, the numerosity model explained less variance in the NPC of dyscalculic participants, and individualized network maps assigned less of this cortex to the dorsal attention network and more to the visual-2 network, without equivalent differences in whole-brain network size; a further DAN difference appeared in left lateral frontal cortex. Across participants, perceptual noise correlated with nPRF R², and visuospatial IQ showed a weaker, tentative association with NPC DAN territory. In the following, we discuss the two accounts in turn, then how they relate to each other, and finally what our approach implies for the study of neurodevelopmental differences more generally.

### Domain-specific differences and response caution

Our modelling results suggest that response caution cannot be left out of a mechanistic account of task performance in DD, as it appeared to compensate for, and thereby conceal, an underlying representational deficit. Simpler measures of the magnitude comparison task, such as choice accuracy and the slope of the psychometric curve, showed only mild impairments in the precision of number sense. Once the PMC-RDM accounted for the increased response caution, a robust increase in perceptual noise became apparent. Dyscalculic participants accumulated more evidence before committing to a response, which bought back in accuracy roughly what noisier representations would otherwise have cost, at the price of slower responses^88,89^. Our simulations confirmed this for the present experiment, as a choice-only model fitted to data simulated with increased caution concealed the perceptual noise difference or even reversed its sign. Comparable compensation has been documented in populations with reduced perceptual efficiency. Children with lower mathematical achievement showed reduced drift rates together with greater response caution in magnitude comparison, responding markedly slower with only modestly reduced accuracy^90^, and children in general make the same trade^81–83^. Similar accuracy on a number-sense task is therefore not by itself evidence for normal processing, as it may result from a strategic adjustment that conceals an impairment. In line with this, the Weber fraction in non-symbolic comparison, unlike the drift rate, has been shown to be contaminated by boundary separation, so that more cautious responders appear to have sharper ANS acuity than they do ^91^. The Panamath Weber fraction is subject to the same confound, so the absence of a group difference there provides weaker evidence for intact acuity than it may appear.

These considerations may help reconcile a divided literature, in which some studies argued for perceptual number sense deficits in dyscalculia^10–12^ and others against them^9,92^. If response caution can conceal representational imprecision, both patterns can be expected, depending on how strongly a given task and sample invite this adjustment.

At the neural level, numerosity-tuned responses in the NPC were less pronounced in adolescents with dyscalculia, as indexed by several measures of nPRF-model explained variance. This converges with a prior report of less distinct neural number representations in dyscalculia in a similar age range, based on multivariate pattern analysis^52^. A further measure of neural precision that incorporates a decoding model^44^ was less sensitive to group differences and less related to behavior. The encoding- and decoding-based measures were strongly related but not equivalent (ρ = 0.65), suggesting that they index somewhat different properties of the population response. In particular, the decoding step selects voxels and fits a noise model within each participant, and this individualized fitting may attenuate group differences that remain visible in the whole-mask encoding measures. At this sample size, however, sampling variance remains a further possibility, all the more as the encoding-based difference was the smallest of the neural group effects.

### Domain-general differences and the network allocation of parietal cortex

On the domain-general side, our results align with studies emphasizing the role of visuospatial working memory in dyscalculia and in mathematics more generally. Working memory capacity robustly predicts mathematical achievement and has been described as a source of vulnerability in dyscalculia^9,19,20^, with its relative importance increasing at later stages of learning^93^.

At the neural level, the connectivity results suggest that dyscalculia involves not only less pronounced numerosity tuning in the NPC, but also differences in how this region is embedded in large-scale networks. In dyscalculic participants, the visual-2 network encroached on NPC territory otherwise affiliated with the DAN, suggesting an altered balance of unimodal and transmodal connectivity along the dorsal stream. Visuospatial IQ tended to covary with parietal DAN territory, although this association did not survive correction across all tested brain–behavior pairs, while the amount of visual encroachment showed no behavioral association. The DAN overlaps broadly with what is elsewhere called the multiple-demand system^60,94^, whose relationship to fluid intelligence is well established and classically probed with matrix-reasoning tasks much like the visuospatial subtest used here. Here, the relationship appeared at the level of the network’s parietal patch, which points to the importance of its topology. The NPC sits on the dorsal “where” pathway, assumed to have originally evolved to code relations in physical space^95^ and recently argued to represent one-dimensional relational content more generally^50^. A patch of the multiple-demand network placed there is a plausible location for the mental storage of relational content, metaphorically a computational substrate of the visuospatial sketchpad of working memory^96^. On this reading, a larger patch would offer more computational space for maintenance and integration, and thereby support non-verbal reasoning, consistent with evidence linking fluid intelligence directly to visuospatial working memory^97^. Given the tentative status of the underlying association, this interpretation remains to be tested in larger samples.

Not every neural group difference had a behavioral counterpart. The visual encroachment within the NPC and the smaller left frontal-lateral DAN patch separated the groups but showed no relationship to any behavioral measure we collected. These neural measures may index components involved in dyscalculia for which our test battery provided no handle, or the sample may have afforded too little power to detect such relations after correction across the full matrix. Given its left lateralization, the smaller frontal DAN patch might reflect verbal contributions to mathematics, such as arithmetic fact retrieval^98^ or complex multiplication supported by the phonological loop^99^. Alternatively, it could relate to executive functions and cognitive control, which are important for arithmetic^100,101^ and thought to rely predominantly on prefrontal areas ^102^. Both possibilities point to components of arithmetic cognition that our behavioral measures did not cover.

### How the two accounts relate

Our study focused on two components of mathematical cognition, between which we found a partial dissociation. The brain–behavior link on the domain-specific side was specific to its own neural measure, whereas on the domain-general side both the association and its specificity were less clear. The two neural measures also showed complementary strengths. The network composition of the NPC separated the groups most clearly, whereas the nPRF measure was most closely related to perceptual noise across individuals. As these correlations were computed with group partialled out, a measure can track individual differences closely without separating the groups strongly, and vice versa. At the same time, the slight correlation between the two neural measures, and the fact that they were derived from the same cortex, suggests that the two components are not fully independent. A similar point has been made about arithmetic cognition more broadly, where “weaknesses in one component can ultimately affect performance in others” through complex developmental processes^103^, likely shaped also by emotional factors such as attitudes towards mathematics. Together with the high rates of comorbidity with other developmental disorders ^6,7,104,105^ and the multi-componential nature of mathematics^103^, this suggests that numerical difficulties rarely reflect a single isolated impairment, and that individuals can arrive at the same diagnosis by different routes.

### Implications for studying neurodevelopmental differences

Our finding that parameters of a cognitive model showed group effects that were weak or absent in raw behavioral measures speaks to a more general issue in cognitive neuroscience. Tasks and their summary measures are often implicitly equated with cognitive constructs, although any such mapping depends on assumptions about how the task is performed^106^. Mean accuracy or a Weber fraction is the joint outcome of perception, memory, and the policy governing when to commit to a response. Even paradigms with simultaneously presented arrays, such as the left–right dot comparisons commonly used to assess ANS acuity, require participants to maintain, integrate, and compare two spatially distinct representations. Reduced performance in such tasks should therefore not be attributed to imprecise number sense without modelling the parallel contributions of maintenance and integration. Participants may moreover differ in the strategies they adopt, so that the same performance can be reached in different ways. Computational models make such differences visible, as they describe how the observed choices and response times were generated. In our data, the accuracy bought by a wider threshold was indistinguishable from sharper representations until response times were modelled as well.

An analogous point holds for the neural data. While the NPC was defined by a group-level functional mask, its internal functional organization varied markedly across individuals. Such variability is expected, as association cortices show pronounced individual differences in functional topography that carry behaviorally relevant information^64,65^. Group-average definitions that assume functional alignment, as in activation or connectivity analyses based on average network atlases, may therefore obscure meaningful variability in how networks are distributed within these regions. In both cases, an observation is read as if it indexed one thing, while what produced it has structure that goes unmodelled. Apparent group differences in numerical cognition may thus arise not only from localized representational deficits, but also from differences in how these regions are functionally organized in each individual, a perspective in line with emerging precision neuroscience frameworks^107^.

### Limitations

Several limitations qualify these conclusions. With 33 participants per group, the study was powered to detect moderate-to-large effects only (see Methods). Although no single participant drove the neural group effects, the reported effect sizes may overestimate the true differences, and it remains to be seen how far they generalize to independent and larger samples. One open question in this respect concerns the cognitive state in which connectivity was measured. Individual network maps were derived from connectivity during the magnitude comparison task, and the group-average map showed a markedly larger DAN than the resting-state reference atlas^62^, which suggests that task engagement may recruit additional territory into this network. Dyscalculia-related alterations in visual-system connectivity have been reported during active number comparison^52^ but not at rest^53,56^, so the shift from DAN towards visual-2 territory within the NPC, which is enhanced in DD, might likewise depend on task demands. On the other hand, functional connectivity patterns appear largely stable across states and vary more between than within individuals^108,109^. Whether the network composition differences, and their relation to visuospatial reasoning, are also present in resting-state data or during other numerical tasks therefore remains to be tested.

## Conclusions

The present results add to a growing body of evidence that dyscalculia cannot be adequately explained by a single cognitive deficit or neural locus. Instead, numerical difficulties appear to reflect a broader constellation of interacting cognitive and neural factors, with both numerical representations, (visuospatial) working memory and underlying large-scale network organization playing central roles. This view is consistent with meta-analytic work highlighting modality-specific working memory vulnerabilities in dyscalculia^8^ and with the high prevalence of comorbid conditions such as dyslexia and attention deficit disorders, thus underscoring the complexity and heterogeneity of the disorder^6,7,104,105^.

These considerations together with our cognitive modeling approach argue against studying DD with performance on any single task and equating that with deficits in a unitary cognitive construct^106^. Instead, our results motivate multidimensional, parametric approaches to investigate mathematical learning difficulties^8^. Positioning individuals within a continuous cognitive–neural space, rather than assigning them to discrete diagnostic subtypes, may provide a more faithful account of the mechanisms underlying dyscalculia or other developmental learning disorders.

From a translational perspective, our findings showcase the promise of combining computational with precision neuroscience for improving our understanding of neurocognitive differences between groups of individuals. While the present study was sufficient to detect group-level differences in some cognitive components, future work should study larger samples with extensive behavioral test batteries, cognitive modelling, and deep phenotyping with precision neuroscience approaches, to capture a more holistic and adequate picture of any type of neurodivergence. Such advances may be essential for improving diagnostic stratification and targeted interventions that address the specific cognitive and neural profiles underlying mathematical learning difficulties.

## Methods

### Study setup

#### Participants

33 adolescents and young adults with a diagnosis of developmental dyscalculia (DD) and 33 neurotypical control participants took part in the study. Groups were matched for age and gender (age range: 15–23 years; 28 females per group). Participants with DD had received a formal diagnosis by a licensed professional from the University Children’s Hospital Zurich, Switzerland and were recruited from a pool of former patients who had previously consented to be contacted for research purposes. Only individuals with a documented normal-range intelligence quotient were invited to participate.

Control participants were recruited from the social environment of the DD participants (e.g., best friends) or from secondary-level schools (“Sekundarstufe II”) in and around Zurich. Control participants reported no history of learning disabilities or neurological or psychiatric disorders. All participants had normal or corrected-to-normal vision and no contraindications for MRI. Written informed consent was obtained from all participants prior to participation. The study was approved by the local ethics committee and conducted in accordance with the Declaration of Helsinki. The target sample size was determined a priori by a power analysis based on the difference in approximate number system acuity (Weber fraction) between adolescents with dyscalculia and typically achieving peers reported by Mazzocco et al.^10^ (d = 0.56), the study whose participants were closest in age to ours. With α = 0.05 and [80% power, one-sided], this yielded a target of 82 participants (41 per group). Recruitment ended at 66 participants (33 per group), when the available recruitment channels, including the patient lists of the [University Children’s Hospital Zurich], had been exhausted. To our knowledge, this is nonetheless among the largest fMRI samples in dyscalculia research to date. Because the final sample fell short of the target, we additionally report the sensitivity of the achieved design (see Statistical power and robustness).

#### Procedure

Participation involved a single laboratory visit to the Social and Neural Systems (SNS) Lab at the Department of Economics of the University of Zurich, Switzerland and lasted approximately three hours, including scheduled breaks. To reduce participant burden, the study entry questionnaires were completed online prior to the laboratory visit by the adolescent participants (and their parents, where applicable). All remaining assessments were conducted on site.

Upon arrival, participants completed either the MRI session or the behavioral assessment outside the scanner, followed by the remaining part of the study. The order of the MRI session and behavioral assessment was counterbalanced across participants and groups.

Before entering the scanner, participants completed a routine MR safety screening. The scanning procedure and experimental tasks were explained in detail, and participants completed several practice trials of the fMRI task to ensure task comprehension. The MRI session lasted approximately 60 minutes. Participants received a fixed reimbursement of CHF 90 for their participation. In addition, they could earn a performance-dependent monetary bonus, based on their behavior in the magnitude comparison task done in the scanner (payout scheme described below) and in a separate risky decision-making task (CHF 0–120).

#### Magnitude comparison task

Participants performed a sequential non-symbolic magnitude comparison task adapted from Barretto-García and de Hollander et al. (2023)^44^, with task difficulty slightly increased to avoid ceiling effects and optimize behavioral modelling. On each trial, participants judged which of two sequentially presented coin arrays was more numerous. A red fixation cross (1 s) preceded the first array (n1), shown centrally for 600 ms. After a jittered inter-stimulus interval of 6–9 s, during which only the red fixation cross remained on screen, the second array (n2) appeared for 600 ms. At its offset, the fixation cross turned green, cueing the response: participants pressed one of two buttons to indicate whether n1 (left) or n2 (right) had appeared larger, and the chosen number (1 or 2) was briefly displayed as confirmation. Responses could be given until the end of the jittered response phase; button presses made before the cue, or trials without a press inside this window, were classified as missed. No feedback on accuracy or accumulated reward was given during the task. The long inter-stimulus intervals ensured that the haemodynamic responses evoked by the two stimulus presentations could be estimated independently. The task was designed with long inter-stimulus intervals to allow the haemodynamic responses evoked by the two stimulus presentations to be modelled separately in the fMRI data.

The task comprised 180 trials distributed across six runs and lasted approximately 40 minutes. Numerosities in the first array ranged from 5 to 20 items, drawn from a geometric sequence with a common ratio of √2 (5, 7, 10, 14, 20). The numerosity of the second array was derived by multiplying the first-array numerosity by a factor of 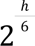 (this factor was adapted compared to the previous study to increase difficulty), with ℎ varying in discrete steps from −6 to 6. Participants were not informed in advance about the range of numerosities presented during the experiment, but familiarized themselves with those already during example trials in the context of the instructions prior to the experiment. Correct responses were associated with a monetary reward of CHF 0.25, although participants were not informed about their ongoing earnings during the experiment.

#### Additional tests and questionnaires

Participants completed a battery of standardized cognitive and self-report assessments outside the scanner. General intelligence was assessed using the IQ screener from the IDS-2 ^72^, which includes subscales for verbal and visuospatial reasoning. Visuospatial short-term memory was assessed using the Corsi block-tapping task ^73^, in which participants reproduce sequences of spatial locations of increasing length; the maximum correctly reproduced sequence length was taken as the Corsi span. Mathematical achievement was assessed using the DEMAT-9 ^74^, a standardized German mathematics test, where points represented problems solved correctly within 3 minutes (max: 50). Math anxiety and math confidence were assessed using self-composed questionnaires inspired by the Mathematikangst-Interview (MAI) ^75^, covering items related to emotional reactions to mathematics and perceived mathematical ability respectively. Approximate number system acuity was assessed using the Panamath task ^76^, which yields a Weber fraction as a measure of the precision of non-symbolic numerosity discrimination.

Unless otherwise noted, all frequentist statistical tests were two-tailed with α = 0.05.

## Data analysis

### Behavioral analysis

Group differences were tested with Mann–Whitney U tests where a measure was not normally distributed (D’Agostino–Pearson test), otherwise with t-tests.

#### Psychophysical model

##### Model specification

As a first behavioral analysis, we used a standard psychophysical (probit) model. We fitted a model where the probability of perceiving n2 as the larger set was a function of an intercept δ and a slope γ times the log-ratio of n2 and n1:

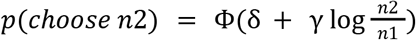

where *n*2 is the size of the second set and *n*1 is the size of the first set.

##### Parameter estimation

We fitted the psychophysical model using the Bayesian hierarchical GLM-estimation package Bambi^110^. As regressors we put in group plus n1 as the base number - to look at potential number range effects (comparing smaller or large numbers, e.g. 5 vs. 7 or 20 vs. 24). Additionally, we also estimated subject-wise random effects on all parameters to obtain subject-wise psychophysical parameters. The following formula was given to the Bambi model estimation procedure:

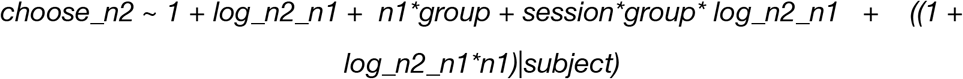

To fit the hierarchical probit model, we used the standard weakly informative priors as implemented by default in Bambi, based on the approach of the well-known *rstanarm* package^110^. Briefly, the priors are Gaussian distributions centered at 0, with as a standard deviation 2.5 times the ratio between the standard deviation of the dependent and independent variable^110^. To sample from the parameter posteriors, we used the no U-Turn sampler (NUTS), which is a self-tuning version of the Hamiltonian MCMC sampler and is particularly efficient for high-dimensional models with correlations between the parameters, such as our model here ^111^. We collected 4 chains of 2000 samples each (1000 burnin). We always visually inspected the traces of the NUTS sampler and made sure the Gelman-Rubin statistic was below 1.05 for all parameters.

### PMC model

Our Perceptual Memory-based Choice (PMC) model assumes that subjects base their choices on noisy neurocognitive representations of the presented non-symbolic magnitudes. All quantities are defined on log-transformed numerosities. On each trial, the representations of the two displays are independent draws

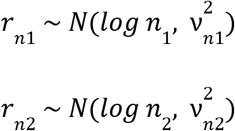

where the noisiness of a representation depends on the order of presentation: because the first display must be retained across the inter-stimulus interval, its noise combines perceptual and memory components in variance,

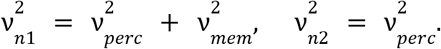

The subject combines each representation with a prior over plausible magnitudes, 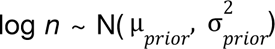, taken to be the objective distribution of presented magnitudes. By standard Gaussian conjugacy, the posterior over the log-magnitude of option *k*, conditional on its representation *r_nk_*, is Gaussian with mean

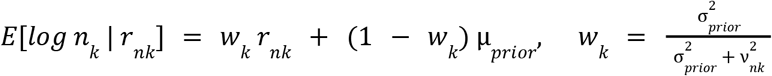

This posterior mean is the subject’s percept of option *k*, and the choice is determined by comparing the two percepts. Conditional on a given representation, the posterior mean is a fixed number; what varies from trial to trial, for the same objective stimuli, is the representation *r_nk_* itself. The trial-to-trial distribution of the percept is therefore obtained by passing the sampling distribution of *r_nk_* through the (affine) posterior-mean mapping, which, conditional on the true numerosities, yields

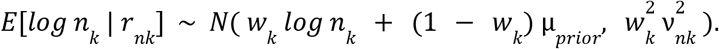

Note that the variance of this distribution, 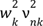, is the marginal variance of the subject’s posterior mean across repeated presentations of the same stimulus. It is distinct from the posterior variance 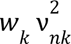, which quantifies the subject’s uncertainty on a single trial but does not itself generate variability in behavior: choices vary across trials because the percept varies, and the percept varies exactly as much as the posterior mean does (cf. the analogous derivation in Khaw, Li, & Woodford (2020)). The mean of the distribution shows the contraction bias directly: each percept is shrunk toward the prior mean by a factor *w_k_*, and more strongly so for the noisier, memory-dependent first option (*w*_1_ < *w*_2_), so that small numerosities are relatively overestimated and large ones relatively underestimated, most of all for *n1*.

Since the two percepts are independent Gaussians, their difference is also Gaussian,

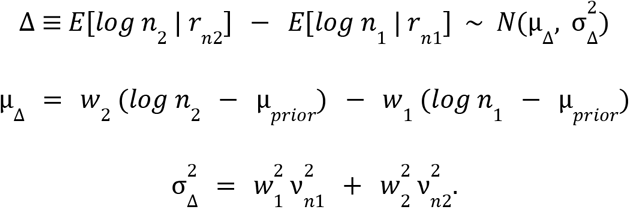

The likelihood of choosing the second option is the probability that this difference is positive:

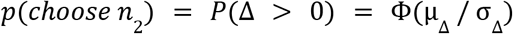

where Φ is the standard cumulative normal distribution. The same marginal variances 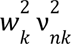 reappear in the dynamic extensions of the model as the (integrated) within-trial diffusion noise of the corresponding accumulators, so that the static and dynamic versions of the model are consistent by construction.

#### Dynamic extension of the PMC model

##### Model estimation

We implemented the PMC, RDM, and PMC-RDM using the Bayesian statistical modeling library pymc 4.3.0 ^v^ ^4.3.0;,112^ and wrapped in our Python package *bauer*^1^. We used a hierarchical (regression) approach for estimation. This means that for all main parameters of the model [ν*_perceptual_*, ν*_memory_*], we assume a group distribution which is a Gaussian distribution. As noises/SDs are necessarily non-negative, we estimated transformed parameters in an unrestricted space [− ∞, ∞] which we then transformed into the non-negative domain [0, ∞] using the softplus function log(1 + exp(*x*)).

For a given parameter θ (ν*_perceptual_* or ν*_memory_*), the subject-session-specific parameter of subject *p* was a linear combination of the mean group parameter µ_θ_ plus an individually-determined offset-parameter δ_θ_ times the standard deviation of the group distribution σ_θ_:

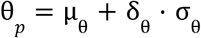

This offset-based specification of the hierarchical model was chosen to prevent the likelihood “funnels” that plague high-dimensional hierarchical models ^113^.

To include potential effects of group affiliation (control versus dyscalculic), we used a regression approach^114^, where a given parameter θ*_p_* (e.g., ν_1_ or σ*_x_*) of subject *p* on a trial *t* (and group *m*) was a linear sum of both an intercept θ_0_ and multiple regression coefficients θ*_n_* times their dummy variable *x*, indicating the group assignment (0 or 1):

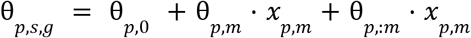

Hence, θ*_p_*_,*m*_ captures any effect related to the group affiliation.

We used mildly informative priors on all group distribution parameters: µ_v_*perceptual*, µ_ν_*memory* ∼ *N*(− 1, 0. 5) (before softplus transformation)

The prior on all group variance parameters was a Half-Cauchy distribution ^115^:

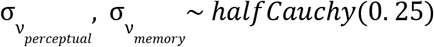

Posterior estimates were obtained using the NUTS-sampler as implemented in pymc 4 with a target acceptance rate of 90%. We obtained 4 chains with 3000 samples each (1500 burnin). If we encountered divergences, we increased the target acceptance rate to 92.5%.

To obtain estimates of the size and presence of an effect, we used “Bayesian” p-values –the probability mass of the posterior estimate above/below 0^116^.

### MRI data

#### Acquisition parameters

We acquired functional MRI data using a Philips Achieva 3T whole-body MR scanner equipped with a 32-channel MR head coil. We collected 6 runs of fMRI data with a T2*-weighted gradient-recalled echo-planar imaging (GR-EPI) sequence (150 volumes + 5 dummies; flip angle 90 degrees; TR = 2286 ms, TE = 30ms; matrix size 96 × 70, FOV 240 × 175mm; in-plane resolution of 2.5 mm; 39 slices with thickness of 2.5 mm and a slice gap of 0.5mm; SENSE acceleration in phase-encoding direction (left-right) with factor 1.5; time-of-acquisition 4:52 minutes). Additionally, we acquired a high-resolution T1-weighted 3D MPRAGE image (FOV: 256 × 256 × 170 mm; resolution 1 mm isotropic; *TI* = 2800 ms; 256 shots, flip angle 8 degrees; *TR* = 8.0 ms; *TE* = 3.7 ms; SENSE acceleration in left-right direction 2; time-of-acquisition 5:35 minutes).

#### MRI preprocessing

Preprocessing on fMRI data was performed using fMRIPrep 20.2.2^117^, which is based on Nipype 1.6.1 ^118^.

##### Anatomical data preprocessing

The T1-weighted (T1w) images were corrected for intensity non-uniformity (INU) with N4BiasFieldCorrection^119^, distributed with ANTs 2.3.3^120^. The T1w-reference was then skull-stripped with a Nipype implementation of the antsBrainExtraction.sh workflow (from ANTs), using OASIS30ANTs as target template. Brain tissue segmentation of cerebrospinal fluid (CSF), white-matter (WM) and gray-matter (GM) was performed on the brain-extracted T1w using fast FSL 5.0.9 ^121^. A T1w-reference map was computed after registration of 3 T1w images (after INU-correction) using mri_robust_template FreeSurfer 6.0.1 ^122^. Brain surfaces were reconstructed using recon-all ^123^, and the brain mask estimated previously was refined with a custom variation of the method to reconcile ANTs-derived and FreeSurfer-derived segmentations of the cortical gray-matter of Mindboggle ^124^.

##### Functional data preprocessing

For each of the 6 fMRI runs per subject, the following preprocessing was performed. First, a reference volume and its skull-stripped version were generated using a custom methodology of fMRIPrep. BOLD runs were slice-time corrected using 3dTshift from AFNI 20160207^125^. Head-motion parameters with respect to the BOLD reference (transformation matrices, and six corresponding rotation and translation parameters) are estimated before any spatiotemporal filtering using mcflirt FSL 5.0.9^FSL^ ^5.0.9;,126^. A B0-nonuniformity map (or fieldmap) was estimated based on two (or more) echo-planar imaging (EPI) references with opposing phase-encoding directions, with 3dQwarp^125^ (AFNI 20160207). Based on the estimated susceptibility distortion, a corrected EPI (echo-planar imaging) reference was calculated for a more accurate co-registration with the anatomical reference. The BOLD reference was then co-registered to the T1w reference using bbregister (FreeSurfer) which implements boundary-based registration^127^. Co-registration was configured with six degrees of freedom. The BOLD time-series (including slice-timing correction when applied) were resampled onto their original, native space by applying a single, composite transform to correct for head-motion and susceptibility distortions. These resampled BOLD time-series will be referred to as preprocessed BOLD in original space, or just preprocessed BOLD. The BOLD time-series were resampled onto the following surfaces (FreeSurfer reconstruction nomenclature): fsaverage, fsnative. Several confounding time-series were calculated based on the preprocessed BOLD: framewise displacement (FD), DVARS and three region-wise global signals. FD was computed using two formulations following Power absolute sum of relative motions ^128^ and Jenkinson (relative root mean square displacement between affines) ^126^. FD and DVARS are calculated for each functional run, both using their implementations in Nipype (following the definitions by Power et al. 2014). The three global signals are extracted within the CSF, the WM, and the whole-brain masks. Additionally, a set of physiological regressors were extracted to allow for component-based noise correction CompCor^129^. Principal components are estimated after high-pass filtering the preprocessed BOLD time-series (using a discrete cosine filter with 128s cut-off) for the two CompCor variants: temporal (tCompCor) and anatomical (aCompCor). tCompCor components are then calculated from the top 2% variable voxels within the brain mask. For aCompCor, three probabilistic masks (CSF, WM and combined CSF+WM) are generated in anatomical space. The implementation differs from that of Behzadi et al. in that instead of eroding the masks by 2 pixels on BOLD space, the aCompCor masks are subtracted from a mask of pixels that likely contain a volume fraction of GM. This mask is obtained by dilating a GM mask extracted from the FreeSurfer’s aseg segmentation, and it ensures components are not extracted from voxels containing a minimal fraction of GM. Finally, these masks are resampled into BOLD space and binarized by thresholding at 0.99 (as in the original implementation). Components are also calculated separately within the WM and CSF masks. For each CompCor decomposition, the k components with the largest singular values are retained, such that the retained components’ time series are sufficient to explain 50 percent of variance across the nuisance mask (CSF, WM, combined, or temporal). The remaining components are dropped from consideration. The head-motion estimates calculated in the correction step were also placed within the corresponding confounds file. The confound time series derived from head motion estimates and global signals were expanded with the inclusion of temporal derivatives and quadratic terms for each^130^. Frames that exceeded a threshold of 0.5 mm FD or DVARS were annotated as motion outliers. Gridded (volumetric) resamplings were performed using antsApplyTransforms (ANTs), configured with Lanczos interpolation to minimize the smoothing effects of other kernels^131^. Non-gridded (surface) resamplings were performed using mri_vol2surf (FreeSurfer). BOLD runs were slice-time corrected using 3dTshift from AFNI 20160207^125^. The BOLD reference was then co-registered to the T1w reference using bbregister (FreeSurfer) which implements boundary-based registration^127^. Co-registration was configured with six degrees of freedom. The BOLD time-series (including slice-timing correction when applied) were resampled onto their original, native space by applying the transforms to correct for head-motion. These resampled BOLD time-series will be referred to as preprocessed BOLD in original space, or just preprocessed BOLD. Many internal operations of fMRIPrep use Nilearn 0.6.2^132^, mostly within the functional processing workflow. For more details of the pipeline, see the section corresponding to workflows in fMRIPrep’s documentation.

#### fMRI analysis - numerical representations

To get a neural surrogate of the noise with which numbers are processed in the brain, we followed earlier work^45,79,133–137^ and used an encoding/decoding-modelling approach, where we inverted an encoding model that describes how a voxel *i* responds to specific stimulus magnitude *s f*(*s*)→*x_i_* in a Bayesian framework, by extending the encoding model to a multivariate likelihood function *p*(*X*|*s*) using a multivariate t-distribution: [*x*_1_,…, *x_n_*] ∼ *f*_1..*n*_(*s*) + ɛ with ɛ ∼ *t*(0, Σ, *d*), where Σ is the residual covariance and *d* is the degrees-of-freedom of the t-distribution. Using an explicit likelihood function allowed us to decode from trial to trial what was the presented numerosity, as well as the fidelity of the neural response. In essence, the model allows us to, for every trial, assess the consistency between the observed BOLD activation pattern and the hypothesized neural representations of numerosity. The fidelity of the neural response was operationalised as the dispersion (standard deviation) of the decoded posterior. Intuitively, the posterior would be less dispersed when the BOLD responses of many voxels agree with a small set of numerosities. When the amplitude of the response across voxels is smaller and/or they are not consistent with the same numerosities, the posterior will be more dispersed, as the decoder is less certain about which stimulus was presented. We postulate, as others have done before^134,135^, that the uncertainty of our decoding of the presented stimulus is related to the fidelity of the *neural representation* of that stimulus. One control participant (sub-03) was excluded from all nPRF analyses because the GLMsingle fit could not be completed for this participant due to a technical problem with the design specification, leaving 65 participants (32 controls, 33 with dyscalculia). This exclusion applies only to the nPRF analyses; the connectivity analyses reported below use the full sample of 66.

##### Single trial estimates

We used the GLMSingle Python package ^138^ to obtain single-trial BOLD estimates. Briefly, the GLMSingle package uses cross-validation to do model selection over GLMs with a) a library of different hemodynamic response functions b) different L2-regularisation parameters to shrink the single trial estimates, combating the issue of correlated single trial regressors ^139^. c) GLMSingle also obtains GLMDenoise ^140^ regressors based on the first *n* PCA components in a set of noise voxels. Noise voxels are defined as having low explained variance in the task-based GLM. The number of PCA components is selected via cross-validation.

As input to GLMSingle, we used both the 24 first and 24 second stimulus presentations per run. For the second stimulus presentations, we modeled all trials with the same numerosity as being in the same condition, to aid GLMSingle with cross-validation (similar numerosities should have similar responses).

##### Numerical receptive field modeling

As a first step, we fitted a numerical receptive field model to all voxels in the brain for each subject separately (over all 180 single trial estimates). Although this method is described in detail elsewhere^26,44^, we briefly describe it here. First, we estimate the mean µ, standard deviation σ, amplitude *A* and baseline *B* of a log-normal receptive field for every voxel in the brain separately, to predict the BOLD response of these voxels to different numerosities.

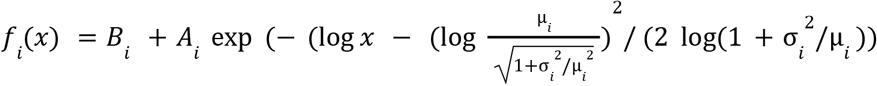

The second part of this equation is a parameterisation of the log-normal probability density function where µ and σ are the mean and standard deviation of the distribution in natural space. Although this parameterisation is somewhat exotic, it is highly useful when plotting the estimated preferred numerosities and their dispersion, for example on the cortical surface.

We fitted the model by first correlating the single trial estimates for the first numerosity stimulus presentation with the predictions of a large grid of 60 µ-s between 5 and 80 and 60 σ-s between 5 and 40. We then estimated *I* and *A* using linear least-squares on the best-correlating µ and σ-parameters. Finally, we used gradient descent^141^ to refine parameters further.

##### Variance explained by nPRF model

To quantify how well the activity within the numerical parietal cortex was described by a numerosity-tuned response, we derived three summaries of each subject’s R² distribution within the group-average NPC mask: its mean, its median, and the number of vertices exceeding an R² of 0.02. Since the model was fitted per subject in individual (volume) space, R² maps were projected onto the *fsaverage5* surface for visualization of cortical maps, and the measures were derived there, so that every subject contributes the same set of vertices. As a robustness check we repeated all three in volumetric space, using the NPC mask warped into each subject’s individual space (Supplementary Table 1).

##### Decoding

To characterize the precision which numerosity can be recovered from the population response we additionally fitted a decoding model in a cross-validate manner as follows. We used a nested cross-validation scheme to select the voxels to decode from for each subject/run similar as described in de Hollander 2024^45^, which avoided the arbitrary choice of the number of voxels within the IPS area. Voxel selection was restricted to individual right numerical parietal cortex (rNPC) masks, obtained by warping a group-average rNPC mask (as in^44^), based on average *R*^2^-maps from the nPRF model described above) from *fsaverage*-surface space to individual surface and then volume (anatomical) space using *neuropythy* (https://github.com/noahbenson/neuropythy).

Specifically, when decoding run *i* (out of a total of 6 runs per session), we fitted the nPRF model on all voxels within the NPC mask on (4-run) subsets of the remaining 5 runs, by leaving another run *j* out *R*__*i*_*j*_. Then, for each of these nested folds *R*__*i*_*j*_, we calculated the *out-of-sample R*^2^ on run *j* for each voxel. Thus, for each voxel and each decoded test run *i*, we had 4 out-of-sample *R*^2^*s*, which we averaged over. For decoding run *i*, we only used voxels that had a mean out-of-sample *R*^2^ larger than 0.

After voxel selection, we used a leave-one-run-out cross validation scheme where the nPRF model was fitted to all runs but the test run, after which also a multivariate noise model was fitted to the residual signals. Specifically, we fitted the following covariance matrix:

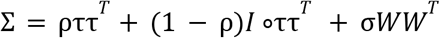

as well as the degrees-of-freedom of a 0-centered, multivariate t-distribution. Briefly, τ is a vector as its length the number of voxels (*n*) in the ROI. It pertains to the standard deviation of the residuals of each voxel. Thus, ρ determines to which extent all voxels correlate with each other (ττ*^T^* is the covariance matrix of perfectly correlated voxels, whereas *I*°ττ*^T^* corresponds to a perfectly diagonal matrix/spherical covariance). *W* is a square matrix of *n* ×*n*. Each element *W_i_*_,*j*_ is the product of the receptive fields of voxels *i* and *j* across stimulus space (*f_i_*(*S*) *f_j_*(*S*)*^T^* with *S* being the entire stimulus space ([5,.., 20] − all the “base” numbers/n1). Thus, the free scalar parameter σ determines to which extent voxels with overlapping receptive fields have more correlated noise.

Once the noise model was fitted, we set up a likelihood function for any multivariate BOLD pattern *X* and for any numerical stimulus *s*:

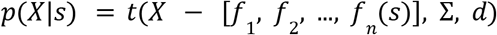

Since we assumed a flat prior on all integers between 5 and 112, we evaluated this likelihood on all these integers and normalized the resulting discretized probability density function (pdf) to integrate to 1. We took the expected value of this pdf to be our estimate of the presented stimulus. Decoding accuracies of individual subjects were then obtained by taking the coefficient of the correlation between those estimates with the true numerosity presented at the respective trials, thus, theoretically ranging from −1 to 1. This was our measure of neural precision, as previous studies have shown that it relates to behavioral precision from numerical tasks^43,44^

#### fMRI analysis - connectivity - precision functional maps (PFM)

To derive individualized network maps, we implemented precision functional mapping pipeline as described in the methods of Gordon et al. (2017)^69^.

##### Post-preprocessing

Functional runs (fMRIPrep-preprocessed, T1w space) were denoised using a Friston 24-parameter motion model ^142^ — six realignment parameters at the current and the preceding TR, together with their squares — plus three physiological nuisance regressors (global signal, white matter, and cerebrospinal fluid, without derivatives or their squares). Frames with framewise displacement greater than 0.2 mm were flagged for scrubbing; rather than being censored outright, flagged frames were first interpolated over using a spectral method, the resulting continuous time series was band-pass filtered (0.009-0.08 Hz), and the flagged frames were then excised (interpolate-then-excise; Power et al. (2014)^143^). Before surface projection, voxels with a temporal coefficient of variation more than 0.5 SD above the local mean (5 mm Gaussian neighbourhood) were masked out to remove dropout and artefact voxels.

##### Surface projection and grayordinate assembly

Denoised volumes were sampled onto each subject’s native midthickness surface using ribbon-constrained mapping (bounded by the smoothwm and pial surfaces), then resampled from native surface space to fsLR 32k standard space (32,492 vertices per hemisphere) via the subject’s FreeSurfer sphere.reg registration and the HCP deformed sphere (adaptive barycentric resampling). Subcortical and cerebellar grey matter (bilateral accumbens, amygdala, caudate, hippocampus, pallidum, putamen, thalamus, and cerebellum), identified from each subject’s T1w-space FreeSurfer parcellation, was retained at the voxel level. Cortical surface and subcortical/cerebellar voxel time series were combined into a single CIFTI dense time series per run and surface-smoothed (sigma = 2.55 mm, geodesic on the surface / Euclidean in the volume), matching Gordon et al. (2017). Runs were concatenated within session after smoothing.

##### Connectivity matrix and thresholding

For each subject, we computed the all-to-all cross-correlation (Pearson) of the concatenated grayordinate time series across all cortical vertices and subcortical/cerebellar voxels. Same-hemisphere node pairs closer than 30 mm were set to zero to reduce the contribution of local spatial autocorrelation to network detection; this 30 mm criterion was approximated as Euclidean distance on midthickness coordinates. We thresholded each subject’s matrix to retain a fixed edge density (the top 0.3%, 0.5%, 1%, 2%, 3%, and 5% of all possible node-pair correlations), with the corresponding r-cutoff for each density approximated per subject (from a random sample of 100,000 node pairs - with ∼85k nodes there are ∼3.6 billion possible pairs).

##### Community detection and labelling

At each density threshold, we ran the Infomap algorithm (two-level, weighted, undirected; ^87^) on the thresholded graph, discarding communities smaller than 400 nodes. Community labels were assigned by matching each community to the reference network with the greatest spatial overlap (plurality vote), using an all-subject average task-connectome network map as the reference; this average map was itself generated by the same pipeline and labelled against the Cole-Anticevic network atlas ^62^. A consensus network map per subject was then obtained by taking, for each node, its label from the sparsest density threshold at which it received a successful reference-matched assignment, with progressively denser thresholds used to fill in nodes that remained unassigned at sparser densities. NPC network composition was quantified, per subject, as the by individual anatomy scaled surface area (cm²) of each canonical network within the NPC mask.

#### DAN patch topology

To characterize the spatial organization of the dorsal attention network (DAN) within and across subjects, we decomposed each subject’s DAN assignment into spatially contiguous sub-regions (”patches”) on the cortical surface. For each subject, vertices labeled DAN in the individual consensus network partition were extracted separately per hemisphere. A vertex adjacency graph was built from the triangulated midthickness surface (each vertex connected to its immediate mesh neighbors, distinct from the correlation-based connectivity graph used for community detection above), and connected-components analysis was applied to the DAN-restricted subgraph to identify spatially contiguous patches. Patch size was quantified as the summed individual anatomical surface area (vertex areas from each subject’s native midthickness surface, converted to cm²). To enable comparison of homologous patches across subjects, we defined a set of reference DAN patches from the group-average network atlas (task-average consensus partition in fsLR 32k). The same connected-components procedure was applied to the group atlas to identify reference patches per hemisphere; patches smaller than 10 vertices were excluded as noise. Each reference patch was assigned an approximate anatomical label (parietal-lateral, frontal-lateral, frontal-medial-dorsal, temporal, frontal-medial, or frontal-insula) based on its centroid coordinates, with the largest patch in each hemisphere/lobe combination receiving the primary label and any additional patches in the same location receiving a numbered suffix. Each subject-specific patch was assigned to the nearest reference patch (within the same hemisphere) by minimum Euclidean distance between patch centroids on the midthickness surface. Where multiple subject patches mapped to the same reference patch, their areas were summed.

#### Brain–behaviour cross-measure correlations

We related eight behavioural/cognitive measures to six neural measures in a single correlation matrix (48 pairs). The behavioural set comprised the three PMC-RDM parameters estimated from the magnitude-judgement task (perceptual noise, memory noise, and boundary separation a; the non-decision time t₀ was excluded) together with five standardized cognitive measures (math skill [DEMAT], visuospatial IQ [Matrizen], verbal IQ [Kategorien nennen], visuospatial working memory [Corsi span], and the log-transformed Weber fraction). Aggregate IQ was not entered separately, as it is a linear combination of the two IQ subscores already in the matrix. The neural set comprised nPRF decodability (r) and mean R², the individual-specific functional-network area of the Visual2 and dorsal-attention networks within the NPC mask, and the area of the two DAN patches that differed between groups (right parietal-lateral, left frontal-lateral).

Associations were quantified as Spearman rank correlations, since several of the measures (network areas, Weber fractions) are bounded and right-skewed, and rank-based coefficients limit the leverage of individual outliers at this sample size. Because both groups are pooled into one matrix, any behavioural and neural measure that each independently differ between dyscalculic and control participants will correlate for that reason alone. We therefore report partial Spearman correlations with group (DD vs. control) entered as a covariate, so that each coefficient reflects covariation over and above what both measures share with the diagnostic split. Each pair was computed on its own complete cases (pairwise deletion; n = 65–66 of N = 66; 33 DD, 33 control). The 48 tests were treated as a single family and corrected using the Benjamini–Hochberg false-discovery-rate procedure; correcting within rows or columns was avoided, as it would understate the size of the search actually performed.

To ask whether a behavioral measure is more closely related to one neural measure than to another, we compared the two dependent correlations using Williams’ test, which accounts for the fact that they share a variable and that the two neural measures are themselves correlated ^144^. These comparisons were computed on group-residualized ranks, so that they test the same partial Spearman associations reported above, and on the cases complete for all three variables. Confidence intervals on the difference between correlations were obtained by bootstrapping over participants (10,000 resamples).

### Statistical power and robustness

Given the modest sample size, we estimated the sensitivity of the design and tested the stability of the main effects. Sensitivity was defined as the smallest effect detectable with 80% power at a two-sided α of 0.05, for the rank-based tests used throughout. Following the asymptotic relative efficiency (ARE) approach, power for the Mann–Whitney U test was computed from the noncentral t distribution with the sample size scaled by its ARE relative to the t-test under normality (3/π; 33 vs. 33 participants), and power for partial Spearman correlations from the Fisher z approximation with the sample size scaled by the ARE of Spearman’s relative to Pearson’s correlation (9/π²; n = 65, one covariate). With 33 participants per group, the design could thus reliably detect group differences of d = 0.72 (0.76 in the least favourable case, ARE = 0.864) and partial correlations of |ρ| = 0.36, so that smaller effects may have gone undetected and the sizes of detected effects may be overestimated.

Effect sizes for group differences are given as Cohen’s d, with 95% confidence intervals obtained by bootstrapping within groups (5,000 resamples). To test whether single participants drove the results, we repeated each neural group comparison, and the brain–behavior correlations shown in Fig. 4B and C, leaving out one participant at a time, using the same test as in the full sample. For the correlations, the full matrix and its FDR correction were recomputed in each leave-one-out sample. We report the range of the resulting statistics and the number of leave-one-out samples in which an effect remained below α = 0.05. These analyses did not include the hierarchical Bayesian behavioral model.

## Footnotes

1 https://github.com/ruffgroup/bauer/

