## Supplementart Material for "Domain-specific and domain-general signatures of dyscalculia in computational models of behavior and parietal cortex"

Supplements

**Supplementary note 1. Relations between the three levels of description of the magnitude comparison task.** To check whether the model parameters captured what they were meant to capture, we tested how the three levels of description of the task (raw performance, the probit model, and the PMC-RDM) related to one another across participants (Supplementary Figure 1). As expected, response time correlated with boundary separation (ρ = 0.51, p(FDR) < 0.001), non-decision time (ρ = 0.38, p(FDR) < 0.01) and perceptual noise (ρ = 0.34, p(FDR) < 0.05), all three of which are meant to influence how long a decision takes. Also as expected, perceptual noise was negatively related to the probit slope (ρ = −0.72, p(FDR) < 0.001), which in turn related to accuracy (ρ = 0.69, p(FDR) < 0.001), establishing the link between these different measures of behavioral sensitivity. Memory noise was most closely tied to the number-range effect of the probit model (ρ = 0.55, p(FDR) < 0.001), as expected if the first stimulus is attracted more strongly toward the prior while it is held in memory. Accuracy, which on its own cannot identify where imprecision arises, correlated equally strongly with the two noise parameters (both ρ = −0.47, p(FDR) < 0.001). Within the PMC-RDM itself, perceptual and memory noise correlated across participants (ρ = 0.43, p(FDR) = 0.001), indicating that they are distinguishable but not fully orthogonal. There was also a weak indication that participants with noisier perceptual estimates adopted a higher decision threshold (ρ = 0.27, p = 0.032, p(FDR) = 0.082), as would be expected if response caution is increased to compensate for imprecise representations.

**Supplementary table 1.** Robustness checks on nPRF R2 measure group comparisons. A) Two types of measures for different ranges: ‘number of vertices’ above the R² threshold at four thresholds (0.01–0.05) and ‘mean R²‘ of the top-ranked vertices at four fixed percentages of the NPC mask (2/5/10/20%, i.e. n = 30/76/151/302 of 1510 vertices), all in *fsaverage5* B) with estimates derived in native volumetric (T1) space and NPC mask.

A)

| **measure** | **Control (mean ± SD)** | **Dyscalculia (mean ± SD)** | **Statistic** | **p** |
| --- | --- | --- | --- | --- |
| **n_vertices(r2>0.01)** | 1330.5 ± 84.3 | 1242.1 ± 141.8 | U = 720.50 | 0.012 |
| **n_vertices(r2>0.02)** | 664.9 ± 244.9 | 513.8 ± 249.3 | t = 2.46 | 0.016 |
| **n_vertices(r2>0.03)** | 325.8 ± 227.6 | 216.4 ± 195.7 | U = 700.00 | 0.024 |
| **n_vertices(r2>0.05)** | 104.2 ± 120.4 | 58.6 ± 95.3 | U = 711.00 | 0.017 |
| **mean_R2(top2%)** | 0.0742 ± 0.0289 | 0.0609 ± 0.0282 | U = 709.00 | 0.018 |
| **mean_R2(top5%)** | 0.0632 ± 0.0241 | 0.0522 ± 0.0238 | U = 712.00 | 0.016 |
| **mean_R2(top10%)** | 0.0545 ± 0.0206 | 0.0452 ± 0.0201 | U = 703.00 | 0.022 |
| **mean_R2(top20%)** | 0.0454 ± 0.0167 | 0.0378 ± 0.0159 | U = 700.00 | 0.024 |

B)

| **measure** | **Control (mean ± SD)** | **Dyscalculia (mean ± SD)** | **Statistic** | **p** |
| --- | --- | --- | --- | --- |
| **mean_R2** | 0.0242 ± 0.0075 | 0.0203 ± 0.0071 | U = 674.00 | 0.014 |
| **median_R2** | 0.0166 ± 0.0044 | 0.0142 ± 0.0039 | U = 684.00 | 0.009 |
| **area(r2>0.02)** | 616.7 ± 150.3 | 529.5 ± 158.3 | t = 2.24 | 0.029 |

**Supplementary table 2.** Whole-brain functional network surface area by group. Values are reported as mean ± standard deviation. Group differences were assessed using independent-samples t-tests or Mann–Whitney U tests depending on normality. P-values were corrected for multiple comparisons using bonferroni correction. Only networks are listed which were detected in every subject

| Network | Control (cm² ± SD) | Dyscalculia (cm² ± SD) | Statistic | p (raw) | p (FDR) |
| --- | --- | --- | --- | --- | --- |
| Visual2 | 276.28 ± 39.33 | 278.60 ± 35.11 | t = -0.25 | 0.801 | 1.000 |
| Somatomotor | 213.39 ± 42.15 | 210.23 ± 37.69 | t = 0.32 | 0.749 | 1.000 |
| Cingulo-Opercular | 256.95 ± 50.89 | 225.41 ± 52.20 | t = 2.49 | 0.016 | 0.109 |
| Dorsal-attention | 260.38 ± 54.25 | 235.75 ± 56.05 | t = 1.81 | 0.074 | 0.521 |
| Frontoparietal | 66.45 ± 47.92 | 89.05 ± 56.85 | t = -1.70 | 0.095 | 0.665 |
| Auditory | 69.22 ± 23.41 | 69.79 ± 22.83 | t = -0.10 | 0.921 | 1.000 |
| Default | 509.65 ± 59.46 | 495.37 ± 53.98 | t = 1.02 | 0.311 | 1.000 |

**Supplementary table 3.** Group comparison of individual anatomical patch area for the eight bilateral dorsal attention network reference patches, sorted by raw p-value. Group sizes varied by patch (control 30--33, dyscalculia 28--33), since not every reference patch was recoverable in every participant's individual network partition.

| Patch | Control (cm² ± SD) | Dyscalculia (cm² ± SD) | Statistic | p (raw) | p (FDR) |
| --- | --- | --- | --- | --- | --- |
| L frontal-lateral | 36.02 ± 14.69 | 25.04 ± 16.42 | U = 804 | <0.001 | 0.002 |
| R parietal-lateral | 62.18 ± 14.51 | 51.89 ± 11.39 | U = 769 | 0.004 | 0.016 |
| L parietal-lateral | 55.82 ± 13.61 | 49.05 ± 11.48 | U = 718 | 0.027 | 0.071 |
| L temporal | 11.93 ± 6.08 | 14.46 ± 7.52 | U = 443 | 0.195 | 0.390 |
| R frontal-lateral | 41.91 ± 21.78 | 36.10 ± 22.05 | t = 1.05 | 0.297 | 0.475 |
| L frontal-medial-dorsal | 7.44 ± 5.78 | 9.17 ± 7.09 | U = 491 | 0.497 | 0.662 |
| R temporal | 12.45 ± 6.73 | 13.24 ± 7.82 | t = -0.43 | 0.670 | 0.766 |
| R frontal-medial-dorsal | 7.20 ± 6.89 | 7.20 ± 6.20 | U = 409 | 0.870 | 0.870 |

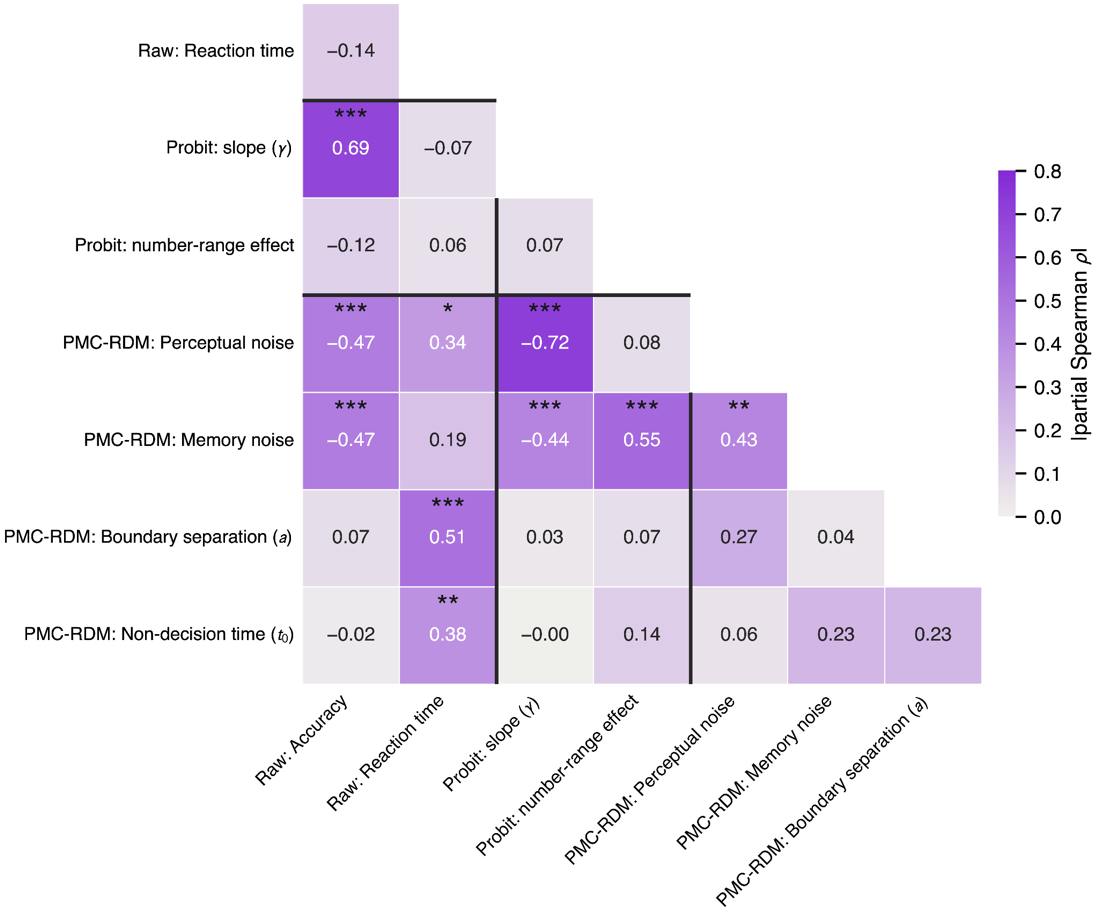

**Supplementary Figure 1.** Associations among the measures derived from the magnitude comparison task. Each cell gives the partial Spearman correlation between two measures, with group (dyscalculia vs.\ control) entered as a covariate, so that a coefficient reflects covariation over and above what both measures share with the diagnostic split. Printed values keep their sign; cell shading encodes |ρ|. Asterisks mark cells surviving correction for multiple comparisons (Benjamini--Hochberg FDR; *** p < 0.001, ** p < 0.01, * p < 0.05). Correction is applied over the 28 unique combinations (half of the symmetric matrix). Rules separate the three levels of description of the same task: raw performance (accuracy, reaction time), the psychophysical probit model (slope γ, number-range effect), and the PMC-RDM (perceptual noise, memory noise, boundary separation a, non-decision time t₀).

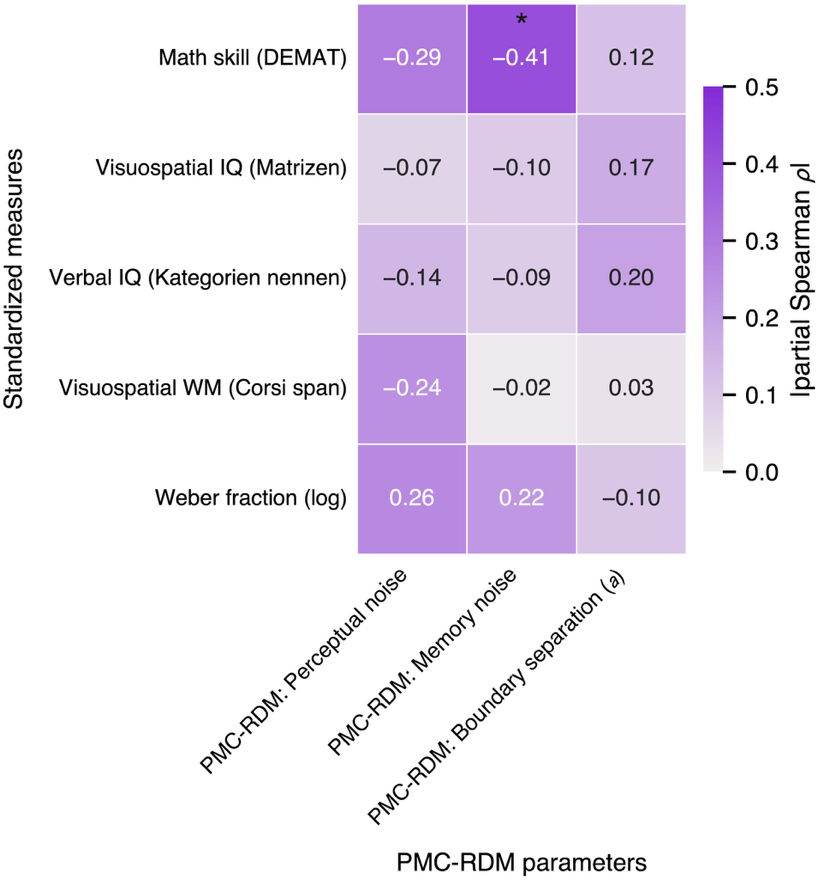

**Supplementary Figure 2.** Associations between the standardized cognitive test results and the PMC-RDM model parameters. Again, correlation values represent partial Spearman correlation with group entered as a covariate and p-values were corrected multiple comparisons (Benjamini--Hochberg FDR; * p < 0.05).

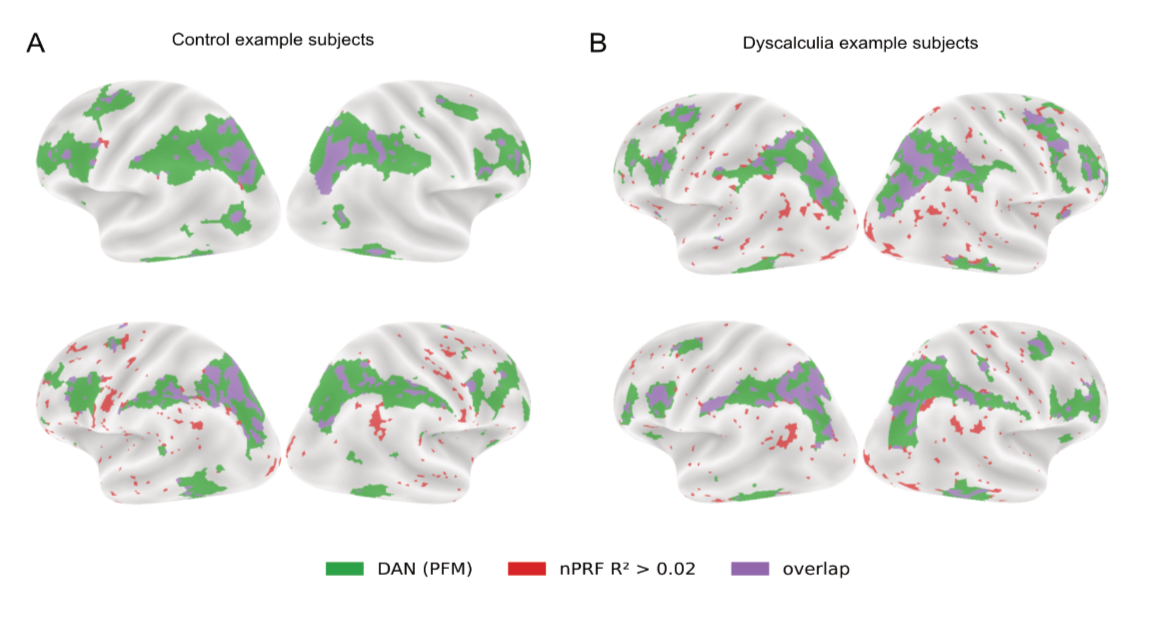

**Supplementary Figure 3.** Example subject overlay brain maps of both models: DAN (PFM) and R2 (nPRF), visualizing how nPRFs are embedded in the parietal DAN patches. Two example subjects from both groups are shown. DAN maps were downsampled to *fsaverage5* space.

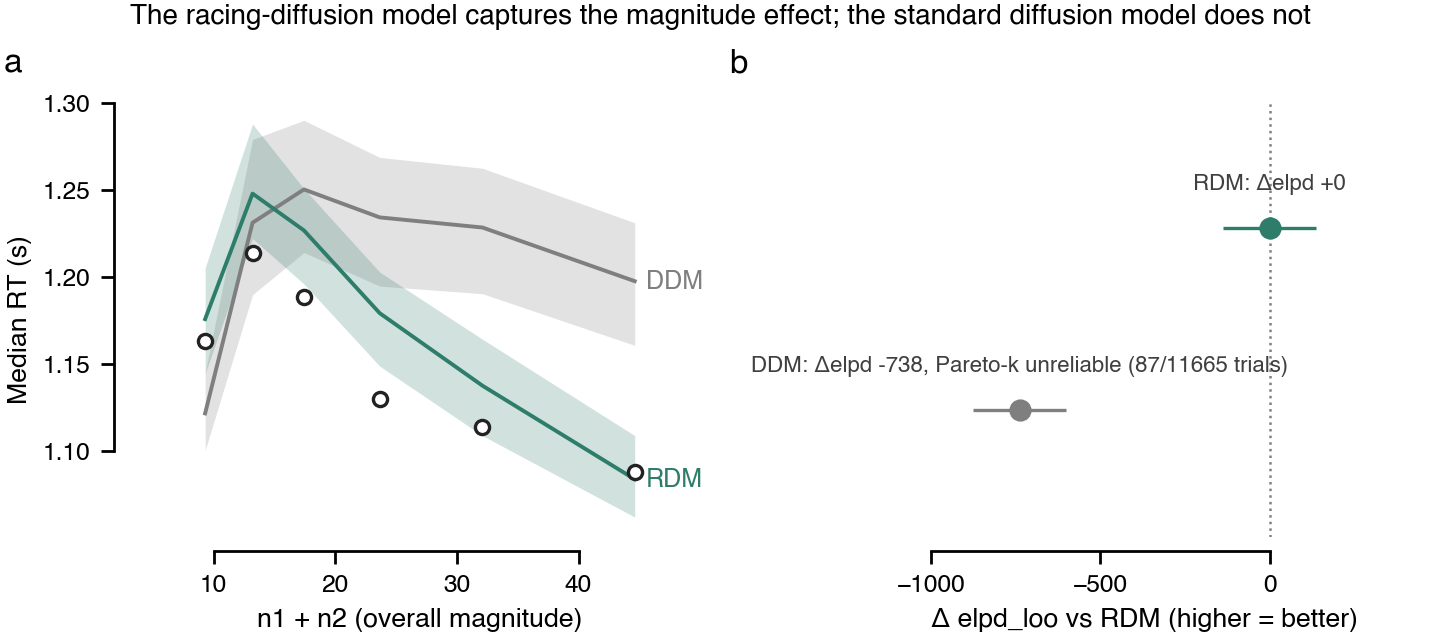

**Supplementary Figure 4. The racing-diffusion model reproduces the overall-magnitude effect on response time; the standard diffusion model does not.** (a) Median response time as a function of overall stimulus magnitude (n1 + n2), pooled across groups (33 control, 33 dyscalculic). Open circles: empirical data. Grey and teal lines: posterior-predictive median RT under the fitted DDM and RDM, respectively (shaded bands: 94% highest-density interval). The RDM reproduces the characteristic decrease in RT with increasing overall magnitude at fixed discriminability; the DDM instead predicts a roughly flat-to-rising RT, because its drift depends only on the evidence difference between the two options, not their sum. (b) Leave-one-out cross-validated model comparison (PSIS-LOO), expressed as the difference in expected log pointwise predictive density (Δelpd_loo) relative to the RDM (dotted line: 0; error bars: ±1 SE). The DDM scores 738 points lower — a decisive difference in the RDM's favour.

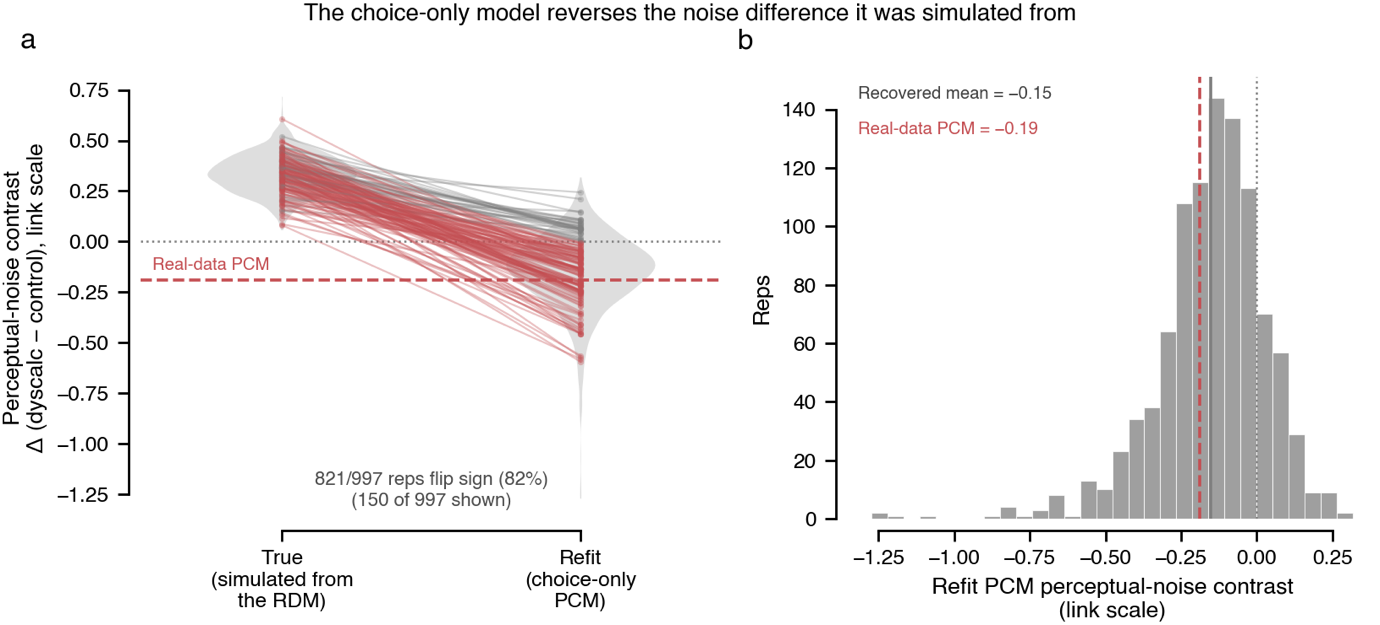

**Supplementary figure 5 A choice-only model reverses the noise difference it was simulated from.** Parametric-bootstrap test of the pure PCM's paradoxical contrast (dyscalculics less noisy, the opposite of the RDM's dyscalculics-noisier result, P = 1.00). If dyscalculics offset extra noise with a higher decision threshold, a model with no threshold parameter should misread the resulting near-normal accuracy as lower noise. Each of 1,000 replicates draws one plausible truth from the RDM (full drift) posterior, in which perceptual noise and threshold are both genuinely higher in dyscalculics, simulates choices and response times for the full 66-subject design, discards the response times, and refits the choice-only PCM to the simulated choices. 997 replicates met the convergence criterion (r̂ ≤ 1.05). (a) Grey violins: distribution of the true simulated contrast (left; positive in 997/997 replicates) and of the refit PCM contrast (right). Thin lines connect 150 randomly chosen replicates, red where the sign flips. (b) All 997 refit contrasts. Grey line, mean −0.15 (95 % CI [−0.16, −0.14]); red dashed line, the real-data PCM estimate (−0.19). Contrasts are dyscalculic − control on the model's unconstrained (pre-softplus) scale; dotted line, 0. The refit reverses sign in 821/997 replicates (82 %, 95 % CI [80, 85]) although the generative noise difference always favoured "dyscalculics noisier", and the real-data estimate falls well inside the simulated distribution (36 % of replicates were at least as extreme). The PCM's paradox is thus expected under the RDM account, and does not indicate a genuine perceptual advantage in dyscalculics. The reversal is specific to the parameter the threshold trades off against: the recovered memory-noise contrast stays positive (dyscalculic > control) in 99.5 % of replicates.
